# Refined cellular activity expression signatures provide a targeted framework to quantify phenotypic intra-tumor heterogeneity in single-cell data

**DOI:** 10.64898/2025.12.19.695441

**Authors:** Angèle Coutant, Lauriane Muller, Cyril Dégletagne, Frédérique Fauvet, Islam El Boudi, Christelle Chassot, Jenny Valladeau-Guilemond, Nathalie Bendriss-Vermare, Anne-Pierre Morel, Maria Ouzounova, Pierre Saintigny, Alain Puisieux, Arnaud Vigneron, Pierre Martinez

**Affiliations:** CASTING - Cancer ecosystem dynamics, adaptation and modeling, Inria, Inserm, Ecole Normale Supérieure de Lyon, Centre Léon Bérard, CNRS, Université Claude Bernard Lyon 1, Lyon, France; Institut Convergence PlasCan, Centre de Recherche en Cancérologie de Lyon (CRCL), INSERM U1052, CNRS UMR 5286, Univ Lyon, Université Claude Bernard Lyon 1, Lyon, France; Plateforme de Génomique des Cancers, Centre de Recherche en Cancérologie de Lyon (CRCL), INSERM U1052, CNRS UMR 5286, Centre Léon Bérard, Université Claude Bernard Lyon 1, Lyon, France; Targeting Non-canonical Protein Functions in Cancer Team, Institut Convergence PlasCan, Centre de Recherche en Cancérologie de Lyon (CRCL), INSERM U1052, CNRS UMR 5286, Centre Léon Bérard, Université Claude Bernard Lyon 1, Lyon, France; Unconventional Tumor Epitopes, Antigen Presentation and T-cell Engineering Team, Institut Convergence PlasCan, Centre de Recherche en Cancérologie de Lyon (CRCL), INSERM U1052, CNRS UMR 5286, Centre Léon Bérard, Université Claude Bernard Lyon 1, Lyon, France; Cancer Immune Surveillance and Therapeutic Targeting Team, Institut Convergence PlasCan, Centre de Recherche en Cancérologie de Lyon (CRCL), INSERM U1052, CNRS UMR 5286, Centre Léon Bérard, Université Claude Bernard Lyon 1, Lyon, France; Gastroenterology and Technologies for Health Team, Centre de Recherche en Cancérologie de Lyon (CRCL), INSERM U1052, CNRS UMR 5286, Centre Léon Bérard, Université Claude Bernard Lyon 1, Lyon, France; Department of Medical Oncology, Centre Léon Bérard, Lyon, France; Institut Curie, PSL Research University, Paris, France; Genetics, Epigenetics and Biology of Sarcomas Team, Institut Convergence PlasCan, Centre de Recherche en Cancérologie de Lyon (CRCL), INSERM U1052, CNRS UMR 5286, Centre Léon Bérard, Université Claude Bernard Lyon 1, Lyon, France; Université Grenoble Alpes, Grenoble, France

## Abstract

Single cell RNA-seq (scRNA-seq) now allows deeper insight into cellular biology at both the individual and population level. Measuring cell-to-cell variations in the population enables quantification of phenotypic heterogeneity in populations in which cell states and identities deviate from healthy transcriptomic profiles. Cellular activities quantifiable using gene set enrichment analyses can provide useful grounds to quantify phenotypic heterogeneity, but the specificity and adequacy of existing molecular signatures for scRNA-seq data is still insufficient. Here we induced 6 activities in vitro, for which we refined existing expression signatures to enhance specificity and detection in scRNA-seq data: epithelial-mesenchymal transition (EMT), DNA repair, responses to interferons α and γ (IFNα and IFNγ, respectively), glycolysis, oxidative phosphorylation (OxPhos). We report new signatures, with much lower redundancy between IFNα and IFNγ, and glycolysis and OxPhos signatures, achieving average AUCs of 0.85 across bootstrapped datasets for each activity. We could use these signatures to quantify phenotypic intra-tumor heterogeneity (ITH) in 20 patient samples and 14 cell lines, observing high correlation with diversity indices in classified healthy cells (p<0.001). Focusing on cancer cells only, we furthermore report higher phenotypic ITH in patients than in cell lines (p<0.001), and in basal tumors (p=0.028).

## INTRODUCTION

Single-cell transcriptomics has revolutionized our ability to characterize RNA expression at the level of individual cells, offering a resolution previously unattainable with bulk methods (1). This has granted deeper insight on the dynamics of RNA expression (2), cell fate decisions (3, 4) and molecular pathways (5, 6). Moreover, single-cell RNA sequencing (scRNA-seq) has facilitated the construction of detailed cellular atlases across tissues, allowing precise quantification of cell (sub)types in both healthy and diseased states (7, 8).

In cancer research, scRNA-seq has been instrumental in dissecting the cellular composition of tumors (9–12), revealing associations between specific cell subtypes and tumor progression or therapeutic resistance. While genetic intra-tumor heterogeneity (ITH) remains a major challenge for diagnosis (13), prognostication (14) and therapy (15), scRNA-seq provides a powerful platform to quantify phenotypic ITH, capturing both genetic and non-genetic sources of variation.

However, as cell subtype classification allows straightforward assessment of the diversity of the surrounding/infiltrating micro-environment (16, 17), quantifying the phenotypic ITH of the cancer cells themselves can prove more challenging. Although cancer cell subtypes have been described indifferent studies (9, 18), they can also represent transient states and may switch over time (19), hampering the biological relevance of classification-based ITH quantification. Furthermore, static classifications can take precedence over more subtle yet functionally-relevant transcriptomic variations (20). To address these limitations, we previously proposed an activity-based framework for quantifying phenotypic ITH, focusing on what cells do rather than what they are, even in absence of well-defined cellular subtypes (21).

Yet, a key challenge remains: the molecular signatures used to infer cellular activities are typically derived from bulk data and are poorly adapted to the sparsity and variability of scRNA-seq (22, 23). In particular, we observed significant redundancy among existing signatures, limiting their specificity and interpretability in single-cell contexts.

To overcome these limitations, we performed in vitro induction of six biologically relevant cellular activities—epithelial-mesenchymal transition (EMT), DNA repair, responses to interferons α and γ (IFNα and IFNγ), glycolysis, and oxidative phosphorylation (OxPhos)—and refined existing gene expression signatures to enhance their specificity for scRNA-seq data. Using AUCell-based enrichment scoring (6), we optimized and validated these signatures across bootstrapped datasets, achieving high performance in distinguishing induced from control cells.

We then applied these refined signatures to public breast cancer scRNA-seq datasets (10, 24), quantifying phenotypic ITH in both patient-derived samples and cell lines. Our activity-based ITH scores correlated strongly with classification-based diversity indices in healthy cells. It furthermore revealed higher phenotypic heterogeneity in patient samples over cell lines, and in basal-like tumors over other subtypes. These findings support the utility of refined activity signatures for robust, classification-independent quantification of phenotypic diversity in cancer.

## MATERIAL AND METHODS

### *In vitro* inductions

EMT and DNA repair activities were induced in pairwise fashion with one negative and one positive control (non-induced and induced cell populations, respectively). To enhance signature specificity for related cellular activities (glycolysis and oxidative phosphorylation; responses to interferons α and γ), the respective inductions were performed in three-way experiments, with one positive control for two negative controls.

#### Epithelial-to-mesenchymal transition (EMT)

MCF10A cells were cultured in Dulbecco’s modified Eagle’s medium (DMEM)/HAMF-12 with 1% glutamax (ThermoFisher Scientific, #31331093), Supplemented with 5% heat inactivated-horse serum (ThermoFisher Scientific, #16050122), 100 U/mL penicillin, 100 µg/mL streptomycin (ThermoFisher Scientific, #15140-122), 10 mg/mL insulin (NovoRapid, NovoNordisk, #7200980), 25 ng/mL Human Epidermal Growth Factor (EGF) (Peprotech, #AF-100-15), 0,5 mg/mL hydrocortisone (Sigma-Aldrich, #H0888) and 100ng/mL cholera toxin (Sigma-Aldrich, #C8052). EMT was induced in MCF10A cells, via the addition of TGF-β in the medium, as in previous work (21, 25). A first analysis was performed after 4 days, when cells started to display mesenchymal morphological features under the microscope (Supplementary Figure 1). Five replicates were furthermore respectively analyzed 1, 2, 3, 4, and 5 days post TGF-β-mediated induction. MCF10A cells unexposed to TGF-β were used as control.

#### DNA repair

MCF10A cells were plated in dishes and grown to 80-90% of confluence before being irradiated at 4 Gy using the Centre Léon Bérard radiology facility. To validate this experimental induction design, cells were lysed four hours post-irradiation in TRIzol™ Reagent (ThermoFisher Scientific, #15596026) following the manufacturer’s instructions to extract RNA. Purified RNA was used for cDNA synthesis and RT-qPCR analysis. Successful induction was controlled by qPCR expression of genes AEN, IER3, IER5, FEN1, PMAIP1 and XPC. RPLP0 expression was used as a housekeeping for qPCR control. Non-irradiated MCF10A cells were used as control (same negative control population as for EMT induction).

#### Responses to interferons α and γ

MDA-MB-468 cells (human mammary adenocarcinoma cells, derived from metastatic site: pleural effusion, DSMZ) were maintained in Leibovitz L15 with 1% GlutaMAX (ThermoFisher Scientific, #31415086) supplemented with 20% FCS (Sigma-Aldrich). In agreement with previous observations of delayed peak response in the interferon γ (IFNγ) case (26), we optimized interferon-response dynamics by exposing MDA-MB-468 cells to interferons α (IFN-α; Schering-Plough) and γ (Peprotech, #200-01A) for 1, 2, and 4 hours, and 4, 10, and 18 hours, respectively. RNA was extracted using TRIzol™ Reagent (ThermoFisher Scientific, #15596026) according to the manufacturer’s protocol and processed for cDNA synthesis and RT-qPCR analysis. Optimal exposure times were determined by monitoring the expression of canonical response genes by qPCR: OAS2 and IRF7 for IFN-α, and CXCL9 and CXCL10 for IFN-γ. MDA-MB-468 cells not exposed to any interferon were used as controls.

#### Glycolysis and oxidative phosphorylation induction

Human mammary epithelial cells (hMEC) cells were grown for 12 weeks in three different culture media based on the same basic medium: DMEM (ThermoFischer Scientific, #A1443001), supplemented with 3% heat-inactivated Fetal Bovine Serum (Eurobio Scientific, #CVFSVF0001), 100 U/mL penicillin, 100 U/mL streptomycin,10 ng/mL EGF. Specific nutrient concentrations were added to the medium to induce each desired condition:

- Neutral medium (control): 10 mg/mL insulin, 0.5 mg/mL hydrocortisone, 1mM pyruvate (ThermoFisher Scientific, #12539059), 6mM glucose (Sigma-Aldrich, #H0888), 1mM L-glutamine (ThermoFisher Scientific, #11500626).
- Glucose-rich medium (glycolysis induction): 20mM glucose, 1mM L-glutamine, no hydrocortisone, no pyruvate, no insulin.
- Glucose-deprived medium (oxidative phosphorylation induction): 0.5 mg/mL hydrocortisone, 1mM pyruvate, 0.5mM glucose, 1mM L-glutamine, 5mM D-galactose (Sigma-Aldrich, #G5388), no insulin.

After the 12-week adaptation period, cells were plated in duplicate in neutral medium. Glucose consumption was monitored by measuring glucose concentration in the cultured medium (supernatant) and protein concentration in cell lysates at 6h and overnight post-plating.

### Glucose consumption assay

#### Protein extraction and quantification

Cells were washed twice with cold phosphate-buffer saline (PBS) and scratched at 4°C in RIPA buffer (50 mM Tris-HCl pH 7.4, 150 mM NaCl, 1% NP-40, 0.5% sodium deoxycholate, 0.1% SDS), supplemented with protease (Roche, #11836145001) and phosphatase inhibitor cocktails (Sigma-Aldrich, #P5726, #P0044). Lysates were sonicated and clarified by centrifugation. Protein concentrations were determined using the Pierce^TM^ BCA Protein Assay Kit (ThermoFisher, #23227), according to the manufacturer’s instructions. Absorbance was measured at 562nm using a microplate reader and concentrations were calculated by interpolating absorbance values from the standard curve generated using serial dilutions of bovine serum albumin.

#### Glucose consumption quantification

Glucose quantification was performed in a 96-well plate using a colorimetric assay. For each well, 5 µL of samples were mixed with 200 µL of reaction mixture and incubated for 1 hour at 37°C. The reaction mixture contained a buffer solution prepared in advance and stored at 4°C, consisting in 0.5mM monobasic potassium phosphate pH 7.4 (Sigma-Aldrich, #P5655), 0.4M sodium hydroxide (Sigma-Aldrich, #S5881), 0.015M Sodium Azide. To this buffer, 50mM 4-hydroxybenzoic acid (Sigma-Aldrich, #H20059), 50mM 4-aminoantipyrine (Sigma-Aldrich, #A4582), 400U/mL Horseradish peroxidase (HRP) (Sigma-Aldrich, #P8375), 4000U/mL glucose oxidase (GOX) (Sigma-Aldrich, #G6125), and distilled water were freshly added. The proportions of each reagent were adjusted so that the final mixture contained 20.2% of buffer, 1% of 4-hydroxybenzoic acid and 4-aminoantipyrine, 0.1% of HRP, 0.19% of GOX and distilled water qsp 200 µL per well. All enzymatic components were added immediately before use to ensure optimal enzyme activity.

After incubation, absorbance was read at 510 nm using a microplate reader (BMG LabTech, Clariostar^®^ Plus). Concentrations were calculated from a standard curve of serial glucose dilution from 0 to 10mM. Glucose consumption was normalized to protein content and averaged over replicates and time points (Supplementary Table 1).

### Primers

All primer sequences are listed in Supplementary Table 2.

### Single-cell RNA-seq data

#### In vitro induction data

Single-cell RNA-seq data was produced in four batches, using the Chromium 3’ kit (10X Genomics), aiming for 1,000 cells per run and 60,000 reads per cell. The first batch included control MCF10A cells, MCF10A cells 4 days post EMT induction, and MCF10A cells 4h post DNA repair induction. The second batch included the control MDA-MB-468 cells, MDA-MB-468 cells post IFN responses to interferons α and γ. The third batch included control hMEC cells, as well as hMEC cells post glycolysis and oxidative phosphorylation induction. Importantly, all populations from the first batch were individually encapsulated, while populations from the following three batches were each pooled before encapsulation using the CellPlex technology (10X Genomics). Cells with fewer than 10,000 or more than 80,000 cells were excluded. Filters on the minimum number of features and maximum percentage of mitochondrial genes acceptable per cell were adapted on a per-sample basis (Supplementary Table 3). Cell-cycle regression was performed using the Seurat package (27, 28), and populations analyzed together were jointly normalized using the SCTransform function.

#### Public data

Two Chromium-based single-cell datasets were downloaded from the GEO database: GSE176078, comprising 20 samples from breast cancer patients (10), and GSE182694, comprising 14 breast cancer cell line samples (24). The scSubtype annotation preferred by the authors (10) was used as patient sample classification, rather than IHC or PAM50. The available processed data from both datasets were integrated and joint-normalized using the Seurat SCTransform function.

### Public RNA expression data

mRNA expression z-scores relative to all samples (log RNA Seq V2 RSEM) data were retrieved from the Firehose Legacy TCGA dataset for breast invasive carcinoma (BRCA), liver hepatocellular carcinoma (LIHC) and kidney renal clear cell carcinoma (KIRC), using cBioPortal (29–31).

### Signature performance evaluation

To ensure that we focused on relevant genes, we based our analysis on “seed” signatures, that have been previously reported in the literature to be related to the 6 selected activities (Supplementary Table 4). We evaluated two types of seed signatures related to each activity: Hallmark gene sets (HM) from the Molecular Signatures DataBase, which were defined on RNA co-expression signal, and Gene Ontology biological processes (GO), which were manually curated from the literature. We further investigated whether the best way to determine whether individual cells presented each activity was to use the entire signatures, or a “refined” versions including only genes considered as significant driver genes for the inactive to active state transition. For each activity, 20 bootstrap replicates were performed by first randomly selecting half from both the positive (post-induction, “active”) and negative (not induced, “inactive”) control populations for discovery purposes. These cells served to first optimize dimensions reduction analyses in semi-automated fashion using the scvelo software (32), then define the significant driver genes. We reduced the number of genes to be analyzed to those from either the HM or GO gene sets, then generated UMAP and velocity graphs with different combinations of values for the n_top_genes (from 25 to the total number of genes in the signature in increments of 25) and min_shared_counts (25, 50, 75, 100) parameters, for the scv.pp.filter_and_normalize function. We selected the optimal combination yielding the most significant difference in mean pseudotime between the inactive (initial state) and active (terminal state) populations (Wilcoxon rank sum test - all tests two-sided unless otherwise specified). Parameters selected for each activity are listed in Supplementary Table 5. From this optimal configuration, we identified driver genes as those with a q-value < 0.05 and a positive correlation with the active state using CellRank (33). Importantly, in each bootstrap iteration, UMAP parameters, pseudotime trajectories, and driver genes were independently redefined within the discovery set, ensuring that, for refined signatures, performance evaluation reflected *ab initio* driver identification rather than parameter reuse. In the EMT case, 4 additional gene sets from Gavish *et al.* (34) were evaluated without refinement, as they were already derived from scRNA-seq data.

For each bootstrap of each activity assessment, the AUCell software (6) was used to score the enrichment for the HM and GO gene sets, both in their entire and refined (significant drivers only) version, on the remaining halves of active and inactive cells. Four different AUCell gene selection thresholds were used: 0.05, 0.10, 0.15 and 0.20, respectively corresponding to including the top 5, 10, 15 and to 20% most expressed genes in the analysis. Performance was then assessed to determine if the AUCell scores could separate the active cells from the inactive ones using Area Under the Curve (AUC) analyses.

To ensure high specificity, IFN responses and metabolic pathways were respectively analyzed in three-way fashion. For instance, for IFNα response, the post-IFNα response induction cells constituted the active population (positive control) while both non-induced and post-IFNγ response induction cells constituted the inactive negative control population, and conversely. For the parameter optimization phase in each bootstrap iteration, all three pairwise population combinations were evaluated, this time selecting the parameter combination yielding the highest sum of W statistic (Wilcoxon rank sum test) over all three pairs, thus maximizing the separation of all 3 populations. These parameters were then used to define the UMAP and velocity graphs for all three populations together (one active, two inactive), then proceed to the driver identification and performance analyses.

### New glycolysis and oxidative phosphorylation signatures

As an alternative to the glycolysis and oxidative phosphorylation HM and GO gene sets, we decided to include the broader “Metabolic process” signature (Gene Ontology GO:0008152) in different combinations with the initial gene sets as possible seed signatures. We evaluated the performance of the refined glycolysis and OxPhos signatures stemming from these combinations based on our *in vitro* induction data, using the same process as for all other activities. Due to difficulties in finding appropriate initial and terminal states in some of the bootstrap replicates, their number was increased to n=40 bootstraps. Replicates in which a trajectory between negative and positive control populations could not be determined were discarded from the evaluation process.

However, the seed signatures being less specific to both activities, we further validated their relevance in two large datasets from cancers known to activate glycolysis (35–37) (Kidney Renal Clear Cell Carcinoma, KIRC) and oxidative phosphorylation (38, 39) (Liver Hepatocellular Carcinoma, LIHC). Driver genes determined via CellRank analyses on the *in vitro* data were cross-validated to select glycolysis drivers upregulated in the KIRC dataset, and to select OxPhos drivers upregulated in the LIHC dataset (p<0.001, t-test). To optimize gene set size, we evaluated in each bootstrap (and for each of the 2 activities) the performance of selecting either the entire list of cross-validated significant drivers, or the top 25, 50, or 75 most significant genes.

### Intra-tumor phenotypic diversity quantification

AUCell scores were calculated for all 6 final activity signatures on all cells from the merged patient and cell lines dataset, using both the optimal AUCell scored determined in the in vitro data and the 0.05 threshold recommended by the authors (6). Activity scores were then linearly normalized so that each activity varied between 0 (lowest score observed in the entire dataset for each activity) and 1 (highest score), and that the maximal and minimal achievable scores for each activity were the same for both cell line and patient samples. As a result, each cell was represented by a vector of 6 values between 0 and 1, one for each of the 6 activities investigated. Pairwise Euclidean distances were calculated between all cancer cells within each population (20 patients, 14 cell lines). The median of each distribution was used to quantify intra-tumor phenotypic diversity. We performed additional analyses only on the non-cancer cells of patient samples, based on the 8 healthy major cell classifications provided by the authors: Endothelial, CAFs, PVL, B-cells, T-cells, Myeloid, Normal Epithelial, Plasmablasts. Our activity-based phenotypic diversity measure was compared to the classification-based Shannon diversity index in each patient’s normal cell contingent. The Shannon diversity index H’ is calculated according to the following formula:

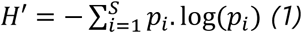

Where *i* is one of all possible subtypes *S*, and *p_i_* is the percentage of subtype *i* in the population.

## RESULTS

### *In vitro* induction of six cellular activities

We first assessed whether existing signatures could be used to accurately predict the presence of six cellular activities in single cell data: epithelial-mesenchymal transition (EMT), DNA repair, responses to interferons α and γ, glycolysis, and oxidative phosphorylation (OxPhos). We used two different and complementary sources as “seed” expression signatures, defined as starting points to optimize the detection of each activity in scRNA-seq data: the MSigDB Hallmark Gene Sets (HM hereafter), based on co-varying RNA expression, and the manually-curated Gene Ontology Biological Process knowledgebase (GO hereafter). Each activity was extrinsically induced *in vitro* in breast cancer cell lines. We evaluated seed signatures either in their entirety or in refined versions based on driver genes, for their ability to distinguish post-induction cells via AUCell (6) enrichment analyses using different gene selection thresholds and a bootstrap procedure (Figure 1, see Methods).

**Figure 1.**
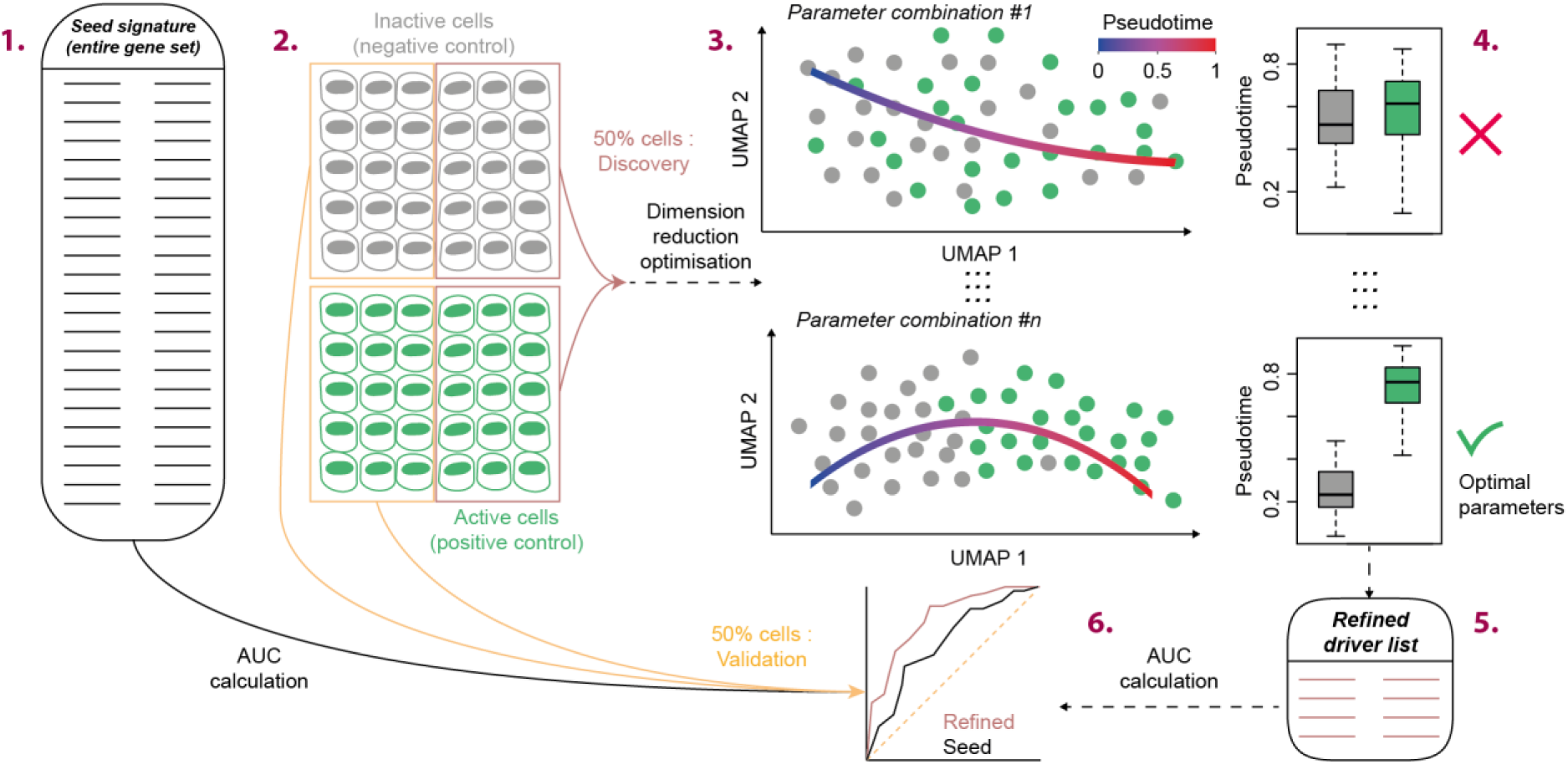
Refined expression signature performance evaluation. Negative (induction-free) and positive (post-induction) control populations were analyzed by scRNA-seq, targeting 1,000 encapsulated cells per population using the following stepwise procedure. **1.** Initial data: a gene set is selected as “seed” signature, either from the MSigDB Hallmark gene sets (HM) or the Gene Ontology biological processes (GO). **2.** Bootstrap: in a bootstrap procedure repeated 20 times, 50% of the positive and 50% of the negative control cells are randomly sampled to create a discovery set (steps 3-5), while the remaining cells form a validation set (step 6). **3.** Parameter space exploration: for each bootstrap, the seed signature is used as initial input for dimensionality reduction analyses using multiple parameter combinations, with the subsequent calculation of the pseudotime underlying the transition from control to post-induction states. **4.** Parameter optimization: the parameter combination yielding the maximum pseudotime separation between control and post-induction cells is selected. **5.** Driver gene identification: driver genes are identified as those whose expression correlated significantly with the transition from control to induced states (correlation > 0; q-value < 0.05), forming refined activity-specific gene sets. **6.** Performance evaluation: signature performance is evaluated using AUC scores on the validation set. The best-performing signature was selected based on the highest median AUC across all bootstraps. When a refined signature outperformed all others during the bootstrap procedure, its final version was determined using the full dataset for parameter optimization and subsequent driver identification.

#### EMT

EMT was induced by addition of TGF-β in the medium of MCF10A cells. Cells were followed over 5 days, with the post-induction state determined after 4 days upon emergence of mesenchymal features observable by microscopy (Supplementary Figure 1). 4 additional EMT signatures from Gavish *et al.* (EMT I to IV) (34), inferred from scRNA-seq data, were also evaluated. The best overall after 4 days was obtained using the EMT II signature (AUC=0.99, 0.2 threshold, Figure 2A-B, Supplementary Figure 2). We however further investigated EMT dynamics through multiple time points, which revealed that signature EMT I better reflected a linear progression between non-induced epithelial cells and nearly fully mesenchymal cells 5 days post-induction (Figure 2C, Supplementary Figure 3). We therefore decided to select the EMT I signature to score the EMT activity in scRNA-seq data.

**Figure 2.**
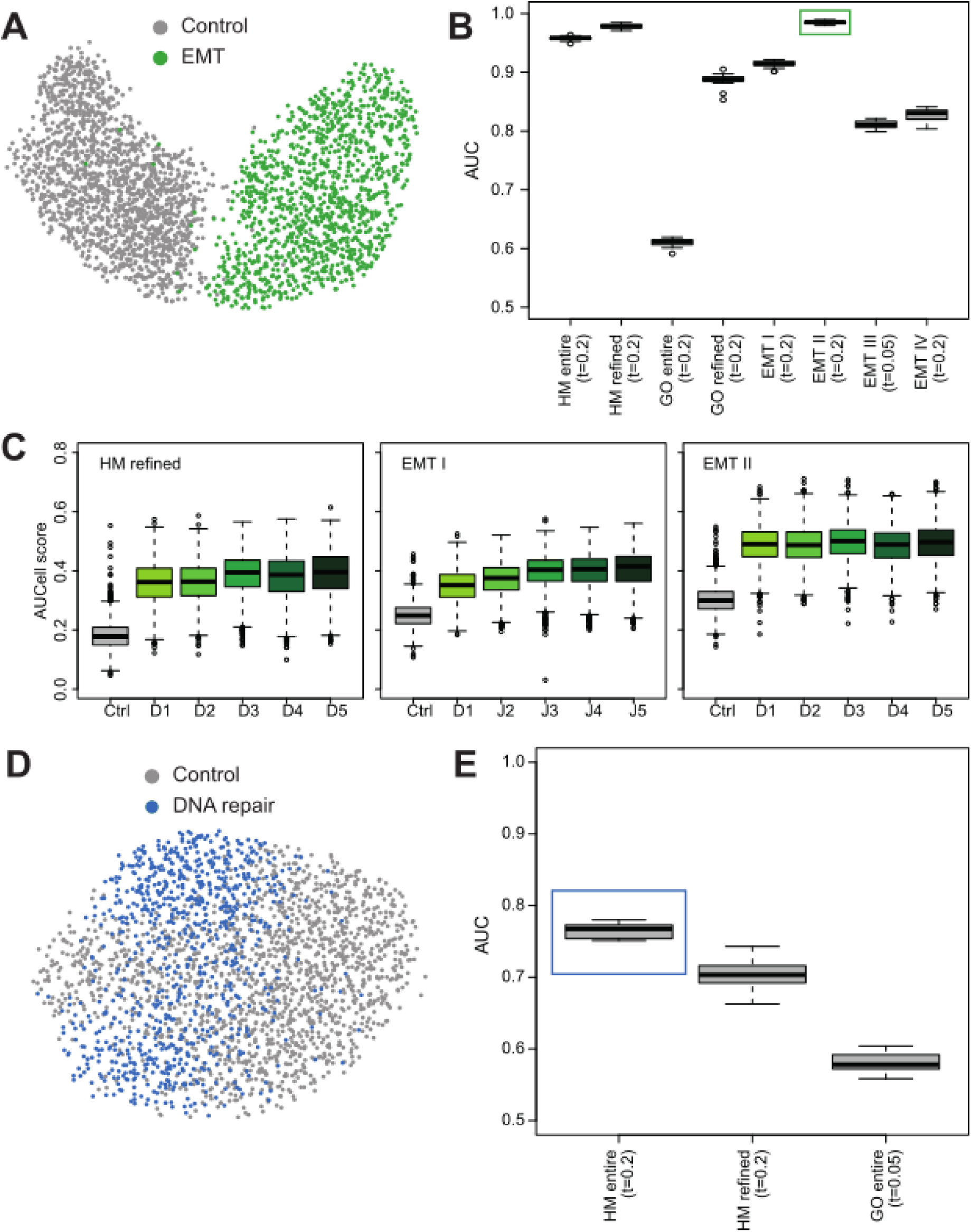
EMT and DNA repair signatures. A) UMAP showing separation of control (grey) and post EMT induction (green) MCF10A cells. B) Best AUC scores across bootstrap iterations, for all EMT gene sets analyzed. The highest AUC is highlighted in green. C) Distribution of enrichment scores over five days post-EMT induction for the EMT HM gene set (left), and EMT I and EMT II signatures (right). D) UMAP showing separation of control (grey) and post DNA repair induction (blue) MCF10A cells. E) Best AUC scores for DNA repair gene sets; the highest AUC is highlighted in blue. Boxplot conventions: boxes represent interquartile range, horizontal lines indicate medians, whiskers extend to 95% CI, and dots denote outliers.

#### DNA repair

DNA repair was induced by irradiating MCF10A cells at 4 Gy, which were analyzed by scRNA-seq 4h later. Irradiated cells displayed increased expression of genes AEN, IER5, PMAIP1 and XPC by qPCR 4h after irradiation (Fold changes > 1.4, Supplementary Figure 4). No expression increase could be observed for genes IER3 and FEN1 (Fold changes >0.8 and <1.2). Drivers could not be determined from the GO gene set, which could thus not be refined and was only evaluated in its entirety. The best performance was obtained using the entire HM signature (Figure 2D-E, median AUC=0.76, 0.2 threshold). Using the standard t=0.05 gene selection threshold, the best AUC=0.68 was in this case obtained using the refined HM signature (Supplementary Figure 5).

#### Responses to interferons α and γ

Responses were induced by incorporating interferon α (IFN α) or γ (IFNγ) into the medium of MDA-MB-468 breast cancer cells, respectively after 4 and 18 hours (Supplementary Figure 6). To enhance the specificity for these related activities, all three populations were analyzed jointly in a three-way design, using 2 negative controls for each activity (Figure 3A, see Methods). We achieved very high performance, with AUCs of 0.97 and 0.99, respectively for IFNα and IFNγ (Figure 3B, 0.2 threshold, Supplementary Figure 7), suggesting that we could define specific signatures adapted to scRNA-seq data for each response. This was confirmed by the low correlation observed between the refined signatures compared to the entire HM and GO gene sets (Figure 3C).

**Figure 3.**
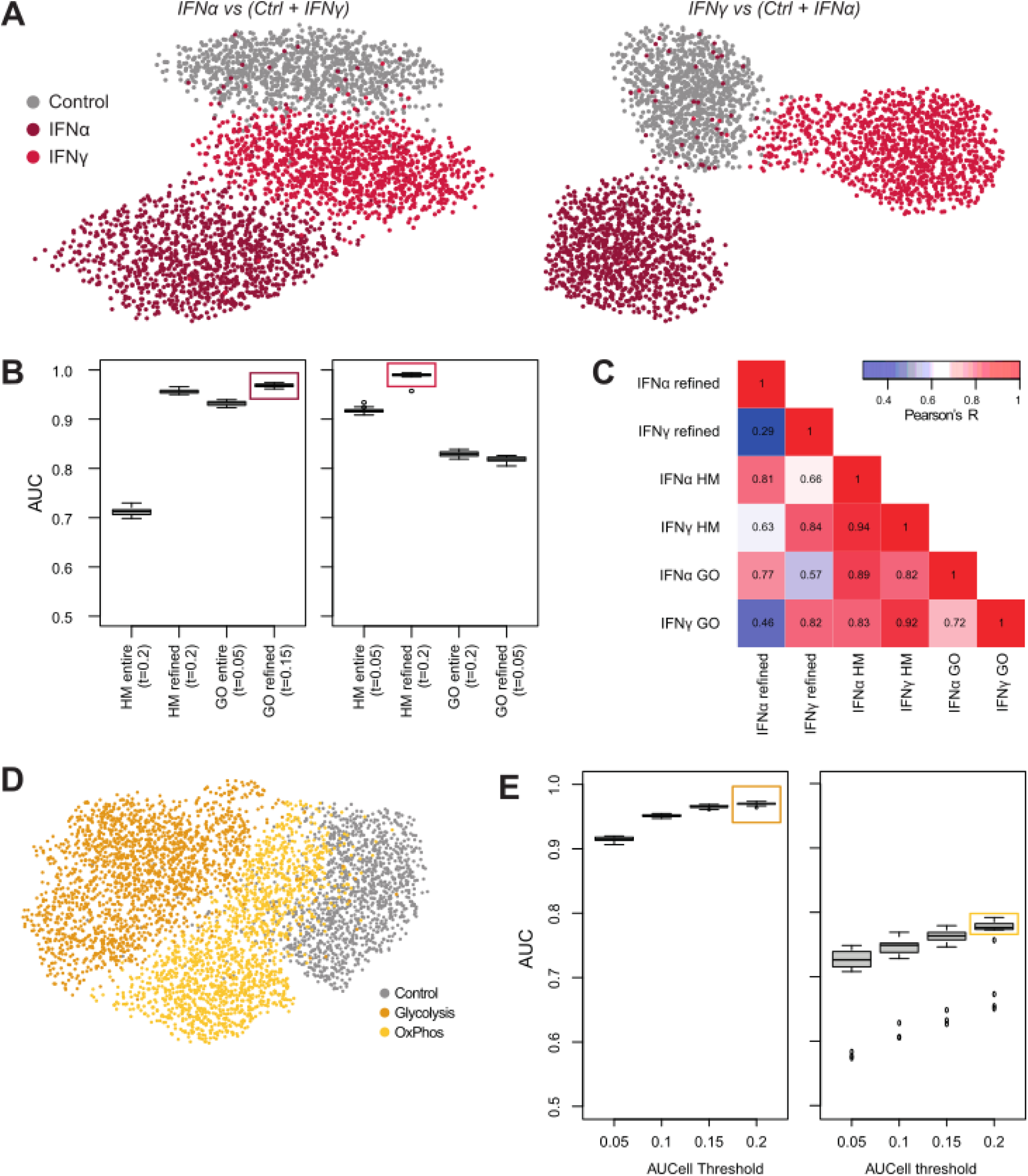
IFN response and metabolic signatures. A) UMAP plots showing separation of control and post-induction populations for IFNα and IFNγ responses (grey, red and pink, respectively). B) Best AUC scores for IFNα (left) and IFNγ (right) gene sets across bootstrap iterations. Best-performing gene sets are highlighted. C) Heatmap showing correlations between full and refined IFN response signatures. D) UMAP plots showing separation of control (grey), glycolysis-induced (orange), and OxPhos-induced (yellow) cells. E) Best AUC scores for glycolysis (left) and OxPhos (right) signatures validated in external datasets. Best-performing gene sets are highlighted.

#### Glycolysis and OxPhos

Both activities were induced in human mammary epithelial cells cultivated in specific nutrient conditions (see Methods). Glucose consumption was highest in the cells previously cultured in the glucose-rich medium, and lowest in the cells previously cultured in the glucose-deprived medium (Supplementary Table 1). Similarly to the interferon responses, these related metabolic activities were analyzed in a three-way design to enhance specificity. However, the HM and GO gene sets could not efficiently separate the three populations, resulting in low AUCs (Supplementary Figure 8). We furthermore observed that the signal from the glycolysis and OxPhos gene sets were largely redundant with each other and proliferation-related signatures in the BRCA breast cancer dataset from TCGA (Supplementary Figure 9).

We thus reanalyzed the cells, using gene set combinations also including the broader “Metabolic process” GO gene set as seed signature (Figure 3D, see Methods). In order to externally validate these gene signatures based on a less specific initial gene set, we further refined them at each bootstrap by selecting the glycolysis drivers most upregulated in clear cell renal cell carcinoma, and the oxidative phosphorylation drivers upregulated in liver hepatocellular carcinoma (TCGA data, see Methods). In our three-way design, these signatures respectively achieved AUCs of 0.97 and 0.78 for the glycolysis and oxidative phosphorylation cellular activities, using respectively 75 and 25 driver genes (Figure 3E, Supplementary Figure 10). These signatures furthermore displayed much lower correlation with each other and proliferation-related signatures in the TCGA BRCA dataset (Supplementary Figure 11).

### Activity-based phenotypic ITH quantification

We scored these 6 activities in two breast cancer datasets, respectively comprising 14 patient and 20 cell line samples, already classified according to the PAM50 signature (Basal, Her2, Luminal A, Luminal B, ER+ and Normal-like, Supplementary Table 6). After normalization to account for inter-signature magnitude variation, we calculated a phenotypic intra-tumor heterogeneity (ITH) score within each sample using pairwise distances (see Methods).

#### Impact of AUCell threshold on phenotypic ITH quantification

We then investigated whether the optimal AUCell thresholds (rather than the standard 0.05 value) defined for each activity using the *in vitro* data were applicable in these new datasets. We found that AUCell scores for each signature did not depend on sequencing depth (UMIs per cell), and that this was not impacted by using optimal thresholds over the standard one (Supplementary Figure 12, p=0.31, t-test). We however found that phenotypic ITH scores were less correlated with UMI counts per cell when using the standard 0.05 threshold (cell lines: Spearman’s ρ=0.13; patient samples: ρ=0.05; Supplementary Figure 13) compared to optimal thresholds (ρ=0.38 and ρ=0.56, respectively), with the latter even showing a significant association in patient samples (p=0.01). In addition, the scores for each activity were more correlated with each other, and therefore more redundant, using optimal thresholds (Figure 4A, Supplementary Figure 14). This suggested that defining optimal thresholds based on our *in vitro* induction data, which were all more relaxed than the standard 0.05 threshold, could induce overfitting and introduce biases in external data analyses. For this reason, we performed all following analyses using the standard 0.05 threshold for signature enrichment analyses using AUCell, and subsequent ITH score calculations. In these conditions, final median AUCs of 0.85 ±13 were achieved on average across the 6 activities, with individual AUCs ranging from 0.68 to 0.97 (Figure 4B). Final gene sets for all 6 activity signatures are listed in Supplementary Table 7.

**Figure 4.**
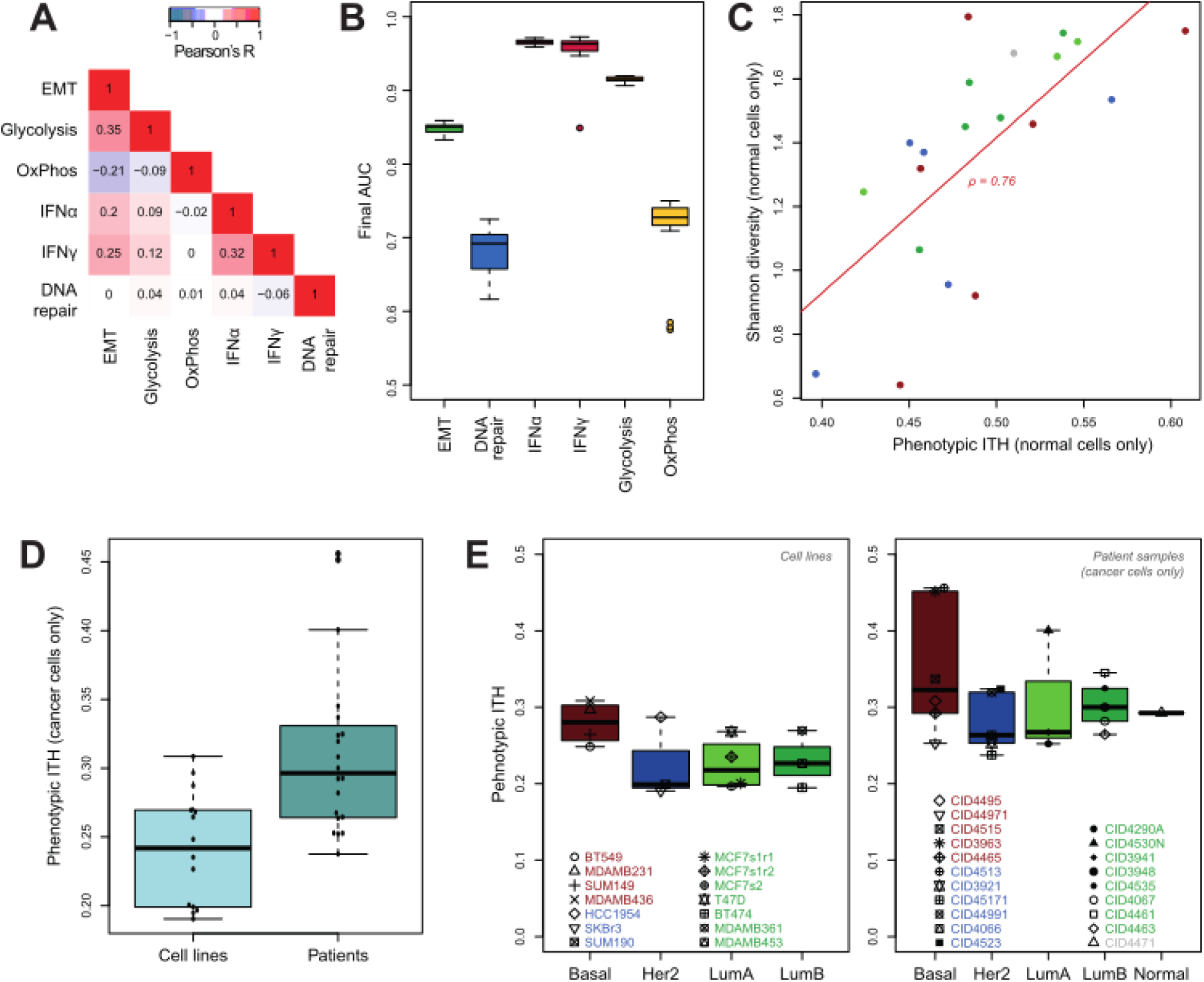
Activity-based phenotypic ITH. A) Heatmap showing pairwise correlations between activity scores (gene selection threshold t=0.05). B) Final AUC scores for each activity signature across 20 bootstrap iterations. C) Correlation between activity-based ITH and Shannon diversity index in normal cells from patient samples. Subtypes are color-coded; red line indicates linear regression fit. D) Distribution of activity-based ITH scores in cancer cells from patient and cell line samples. E) Activity-based ITH scores across breast cancer subtypes in cell lines (left) and patient samples (right).

#### Activity-based ITH reflects classification-based diversity in normal cells

To evaluate the relevance of our activity-based ITH quantification method, we calculated it on the non-cancer cells in the 14 patient samples, and compared it to the Shannon diversity index based on cell type frequencies. We observed a high correlation between the two measures (p<0.001, Spearman’s ρ=0.76, Figure 4C), suggesting that our activity-based phenotypic ITH score accurately reflects phenotypic diversity in populations of heterogeneously classified healthy cells.

#### Refined signatures provide improved quantification compared to entire signatures

We further investigated the impact of using refined signatures rather than the entire MSigDB gene sets for each activity. We found that activity scores calculated using entire gene sets yielded a strong correlation between IFNα and IFNγ response signatures (Supplementary Figure 15A, R=0.92) and worse correlation with the classification-based Shannon diversity index (Supplementary Figure 15B, ρ=0.5). This suggests that our refined signatures helped reduce the redundancy between activities, and increased our accuracy in quantifying phenotypic differences between cells of a population.

#### Basal-like breast cancers display higher phenotypic ITH

Using signatures defined with the standard 0.05 AUCell threshold, we observed that intra-tumor phenotypic diversity was higher in patient samples (Figure 4D, p<0.001, t-test). Although UMI counts per cell were higher in cell lines than in patient samples (Supplementary Figure 16, p<0.001, t test), these were not correlated with activity-based phenotypic ITH in either sample type (Supplementary Figure 13, 0.05 threshold), and thus unlikely to explain the greater ITH of patient samples.

Furthermore, when comparing subtypes, we found that samples associated to a Basal subtype displayed higher phenotypic ITH (Figure 4E, p=0.028, Wilcoxon rank-sum test), particularly in cell line data (p=0.014). This finding was still significant after multivariate linear regression to correct for patient/cell line sample type (p=0.004 without interaction effect, p=0.033 with, Supplementary Tables 8-9). Furthermore, we observed that the diversity of the normal cells in basal tumors was similar to other tumor types, and that phenotypic ITH and TME diversity were not correlated across all patient samples (Spearman’s ρ=−0.01, p=0.977, Supplementary Figure 17). This suggests that the higher ITH observed in basal tumors is more likely due to intrinsic characteristics of basal cancer cells rather than to a superior diversity of the microenvironmental cells with which they interact.

## DISCUSSION

In this study, we refined molecular expression signatures for six cellular activities—epithelial-mesenchymal transition (EMT), DNA repair, responses to interferons α and γ (IFNα and IFNγ), glycolysis, and oxidative phosphorylation (OxPhos)—to improve their specificity and applicability to single-cell RNA-seq (scRNA-seq) data. Starting from existing gene sets, we used in vitro induction experiments to identify driver genes that better reflect activity-specific transcriptional changes in individual cells. This refinement process enabled us to reduce redundancy between related signatures and improve performance in distinguishing post-induction from control populations.

Among the refined signatures, those for IFNα and IFNγ responses achieved particularly high specificity and sensitivity, with minimal overlap between them. This represents a significant improvement over existing bulk-derived gene sets, which often conflate these closely related pathways due to overlapping genes. In contrast, our DNA repair signature showed the lowest performance, which may stem from a mismatch between the specific types of DNA damage induced by ionizing irradiation (40)—such as single- and double-strand breaks, and abasic sites—and the broad scope of a single DNA repair signature encompassing multiple, mechanistically distinct repair pathways. This highlights the challenge of capturing complex, multi-modal biological processes with a single gene set.

Importantly, while area under the curve (AUC) metrics are useful for evaluating binary classification performance, they may not fully capture the biological relevance of continuous transitions, such as those observed during EMT. For this reason, we selected the EMT I signature (34) based on its ability to reflect a gradual phenotypic shift over time, rather than solely on its AUC score. This decision underscores the importance of considering biological dynamics when evaluating signature performance.

By focusing on cellular activity rather than identity, our approach offers a functional lens through which to examine tumor heterogeneity. We used the AUCell software (6) to quantify the enrichment of our refined activity signatures in scRNA-seq data from breast cancer patients and cell lines and leveraged these scores to compute activity-based phenotypic intra-tumor heterogeneity (ITH). Our activity-based ITH scores correlated strongly with classification-based Shannon diversity indices in healthy cells, suggesting that this approach captures biologically meaningful phenotypic variation. Refined signatures furthermore increased this correlation compared to standard ones, suggesting that our signature refinement procedure improved their biological relevance.

Notably, we observed higher phenotypic ITH in patient samples compared to cell lines, which likely reflects the greater variety of interactions cancer cells have with a diverse microenvironment that cannot be fully replicated *in vitro*.

We also found that basal-like breast cancers exhibited higher phenotypic ITH than other subtypes, in both patient and cell line data. This observation aligns with previous reports of increased genomic instability in basal tumors (41), and may help explain their poor prognosis and resistance to therapy (42). Activity-based profiling could thus provide a functional complement to genomic analyses, offering insights into non-genetic mechanisms of tumor adaptation (43) and therapeutic escape.

One limitation of our approach is the sensitivity of AUCell scoring to threshold selection. While relaxing the threshold improved performance in controlled settings, it introduced sequencing depth-related biases in external datasets. This illustrates a trade-off between sensitivity and robustness and the need for further standardization and automatization of activity-based ITH quantification in scRNA-seq data.

Overall, our study demonstrates that activity-based profiling is a promising strategy for quantifying phenotypic ITH in scRNA-seq data. By focusing on cellular function rather than identity, this approach enables classification-independent analysis and reveals biologically relevant patterns of heterogeneity. As more refined scRNA-seq-compatible signatures become available (34), this framework can be expanded and adapted to support deeper functional characterization of tumors and other complex tissues.

## DATA AVAILABILITY

All original data have been deposited as a superseries on the GEO database: GSE310225.

## CODE AVAILABILITY

The code used is deposited on GitHub: https://github.com/AngeleCTNT3/PhenoDiv

## Author contributions

Conceptualization: APM, AP, AV, PM; Data curation: AC, LM, AV, PM; Formal analysis: AC, LM, IEB, AV, PM; Funding acquisition: PM; Investigation: AC, LM, AV, PM; Methodology: AC, LM, CD, FF, CC, JV, NBV, APM, MO, AV, PM; Project administration: PM; Resources: FF, CC, APM, AV, PS, AP; Software: AC, IEB, PM; Supervision: JV, NBV, APM, MO, PS, AP, AV, PM; Validation: AC, LM, FF; Visualization: AC, LM, PM; Writing – original draft: AC, LM, PM; Writing – review & editing: AC, LM, PM.

## FUNDING

This research was financially supported by the Institut Thématique Multi-Organismes (ITMO) Cancer of AVIESAN (Alliance Nationale pour les Sciences de la Vie et de la Santé, National Alliance for Life Sciences & Health) within the framework of the Cancer Plan. AC is funded by a scholarship from the Ligue Contre le Cancer.

## CONFLICT OF INTEREST

AV is a founder and the chief scientific officer of the Sirius NeoSight company, which was not involved in the study design, data collection, analysis, interpretation, or manuscript preparation. Other authors report no conflict of interest.

## Notes

https://www.ncbi.nlm.nih.gov/geo/query/acc.cgi?acc=GSE310225

### Summary of Updates

Author list reordering. Pierre Martinez is corresponding and last author.

