## Supplementary Tables and Figures for "Refined cellular activity expression signatures provide a targeted framework to quantify phenotypic intra-tumor heterogeneity in single-cell data"

**Supplementary Figures**

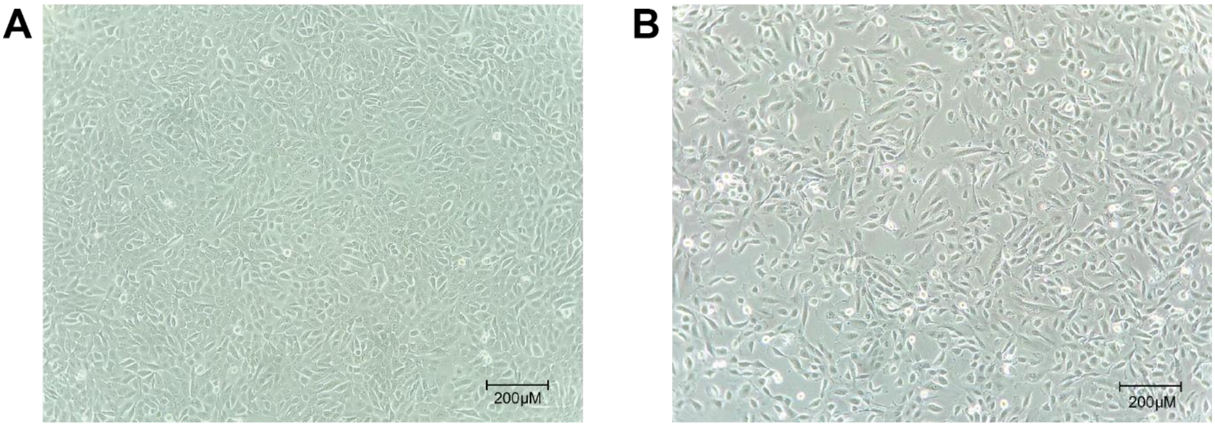


**Supplementary Figure 1 –** EMT induction. A) MCF10A cells in absence of TGF-β. B) MCF10A cells 4 days post TGF-β-mediated EMT induction TGF-β.

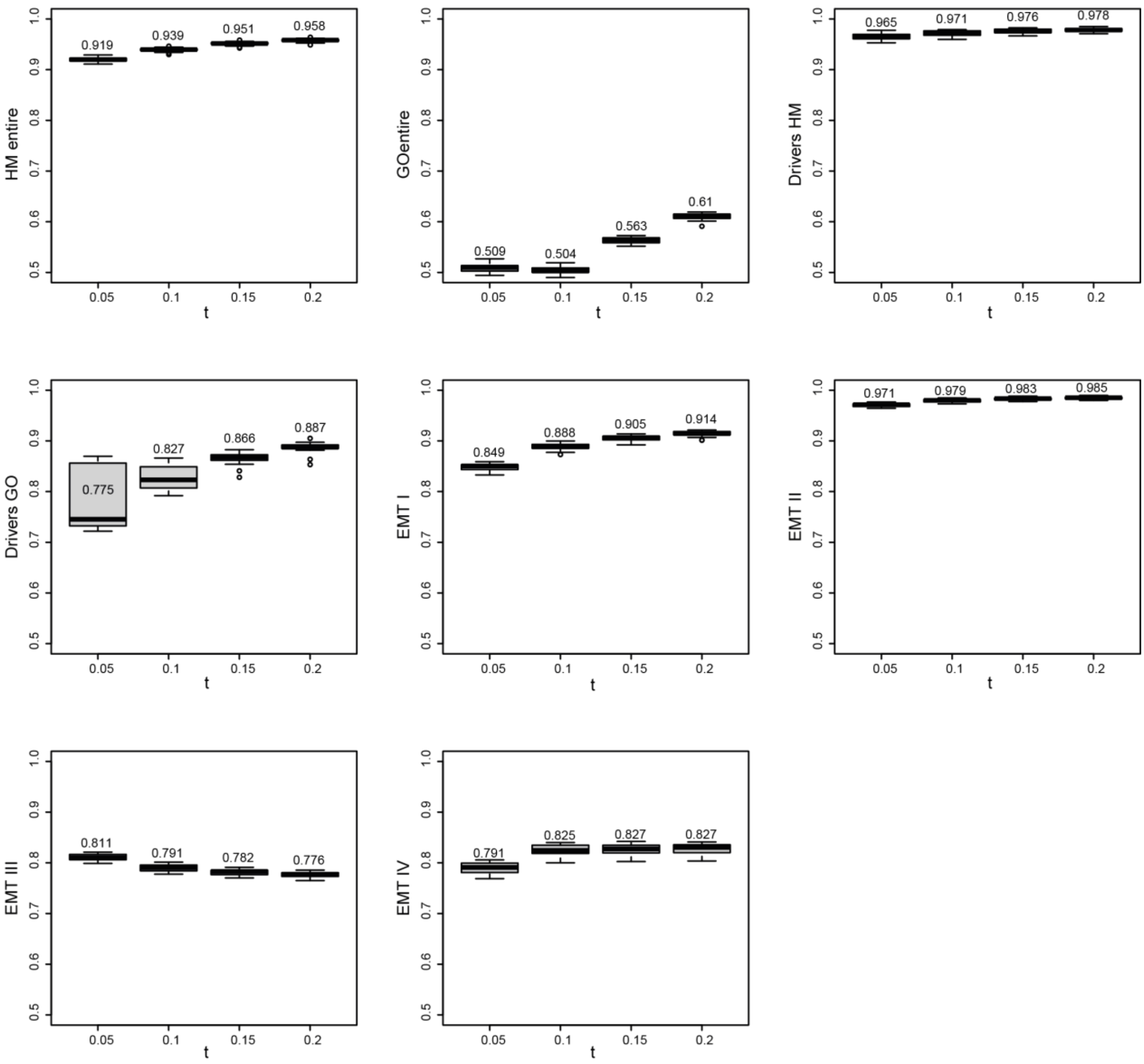


**Supplementary Figure 2** - AUC scores across bootstrap iterations for all EMT gene sets, using 4 different gene selection thresholds (t). Boxplot conventions: boxes represent interquartile range, horizontal lines indicate medians, whiskers extend to 95% CI, and dots denote outliers.

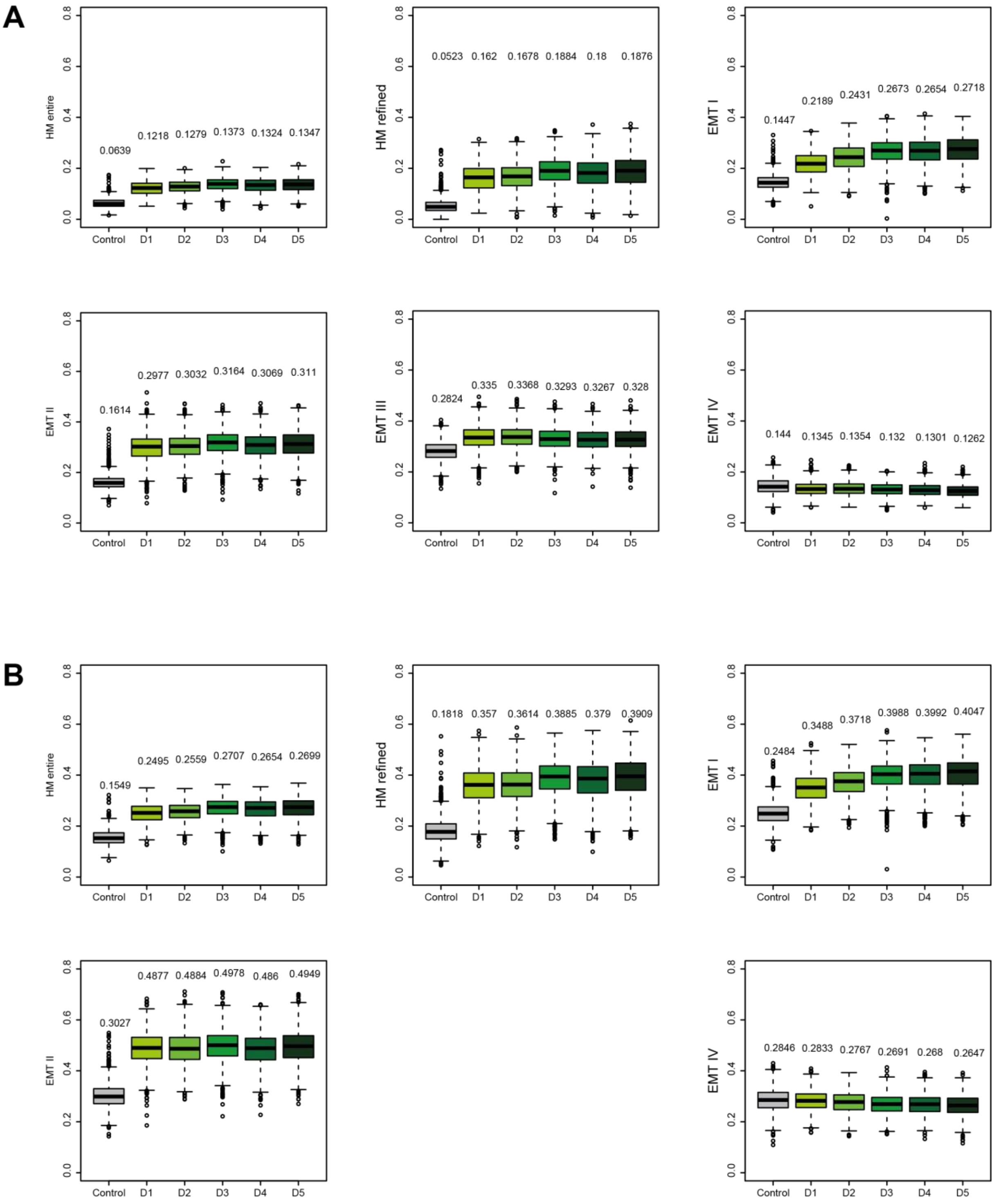


**Supplementary Figure 3** – AUCell scores across bootstrap iterations for all EMT gene sets, using A) the standard 0.05 gene selection threshold t and B) the one providing the highest AUC. The highest AUC was achieved using t=0.2 in all cases but the EMT III gene set (t=0.05, see Supplementary Figure 2).

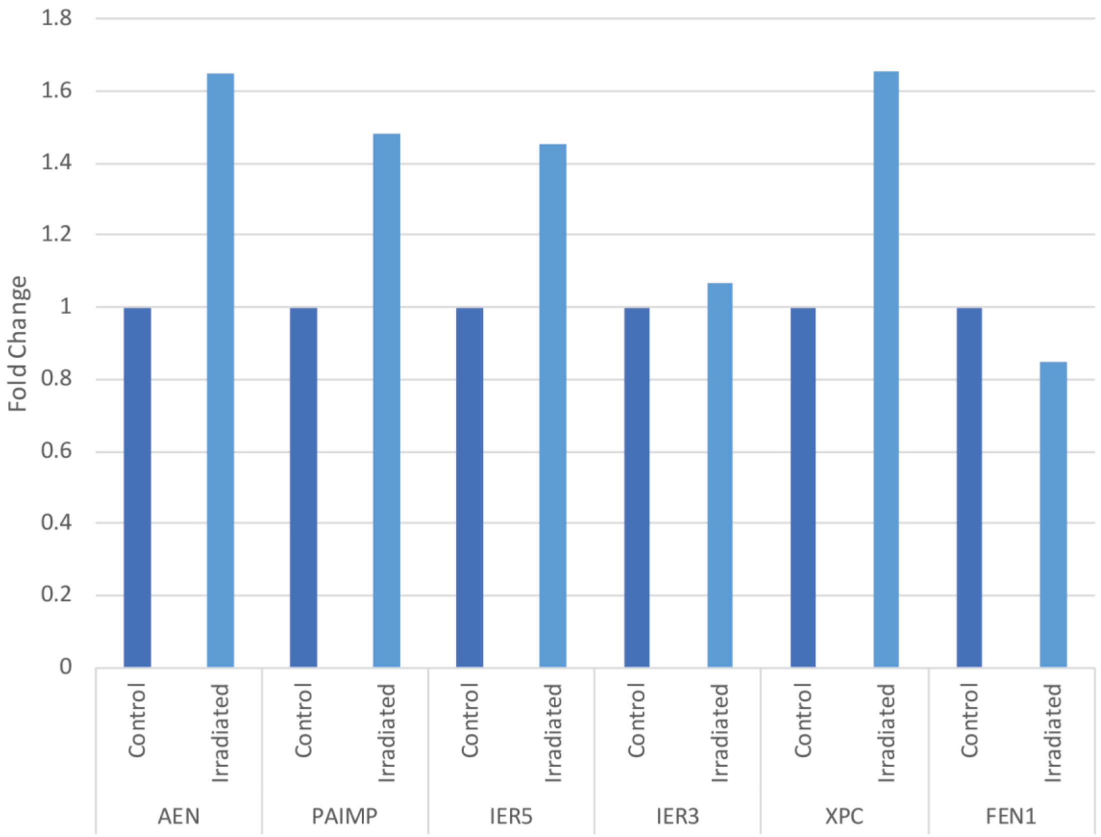


**Supplementary Figure 4** – Gene expression fold change in MCF10A cells 4h after 4Gy irradiation.

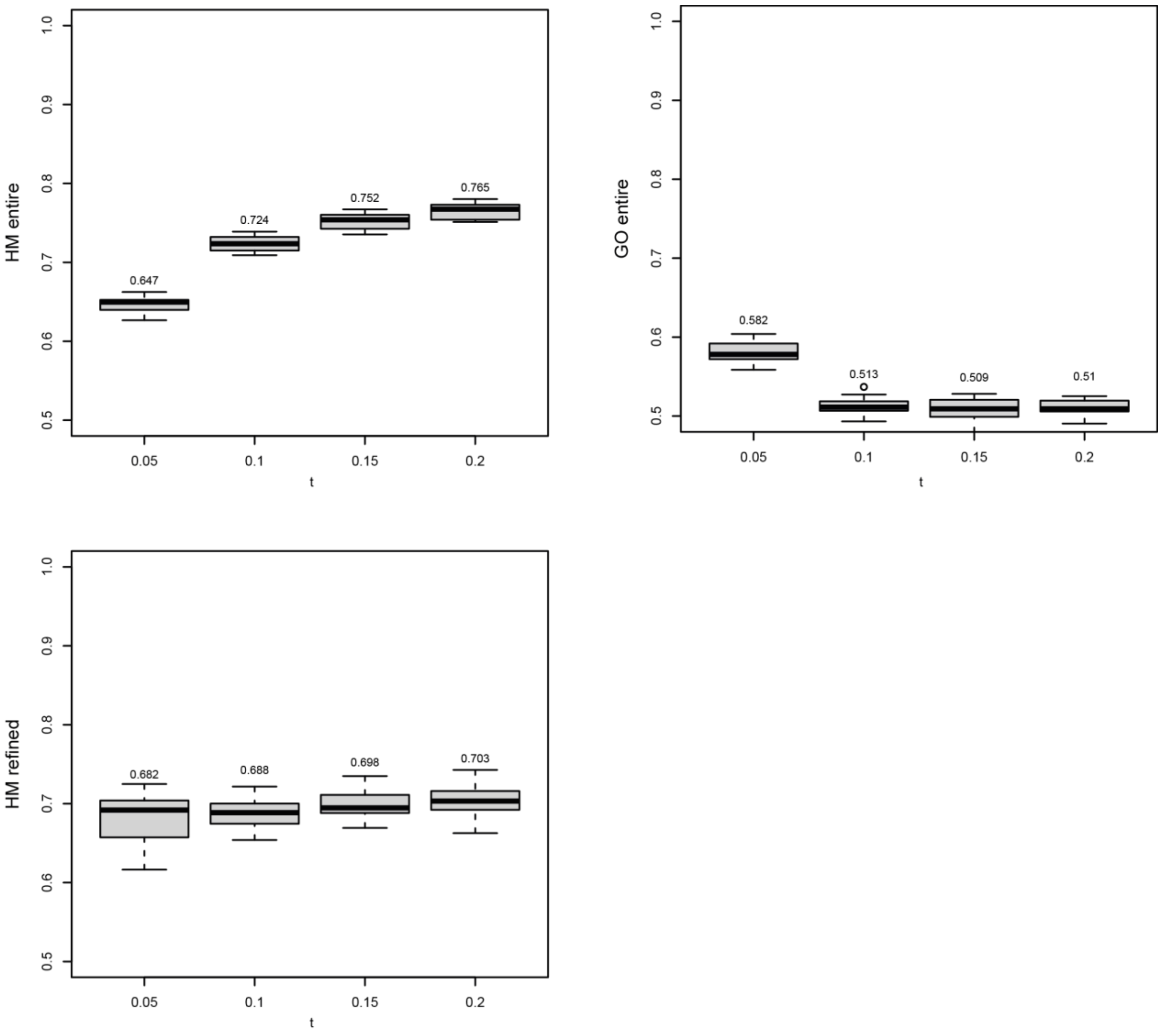


**Supplementary Figure 5** – AUC scores across bootstrap iterations for all DNA repair gene sets, using 4 different gene selection thresholds (t). Drivers could not be identified using the GO “DNA repair” gene set, and thus no refined GO gene set could be evaluated.

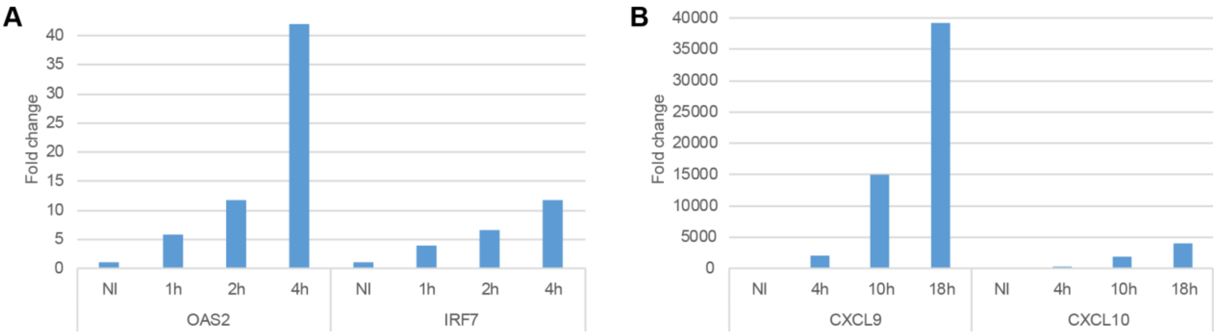


**Supplementary Figure 6** – Gene expression fold change in MDA-MB-468 cells after exposition to interferons A) α and B) λ. Time points on the x-axis, NI stand for non-induced (negative control).

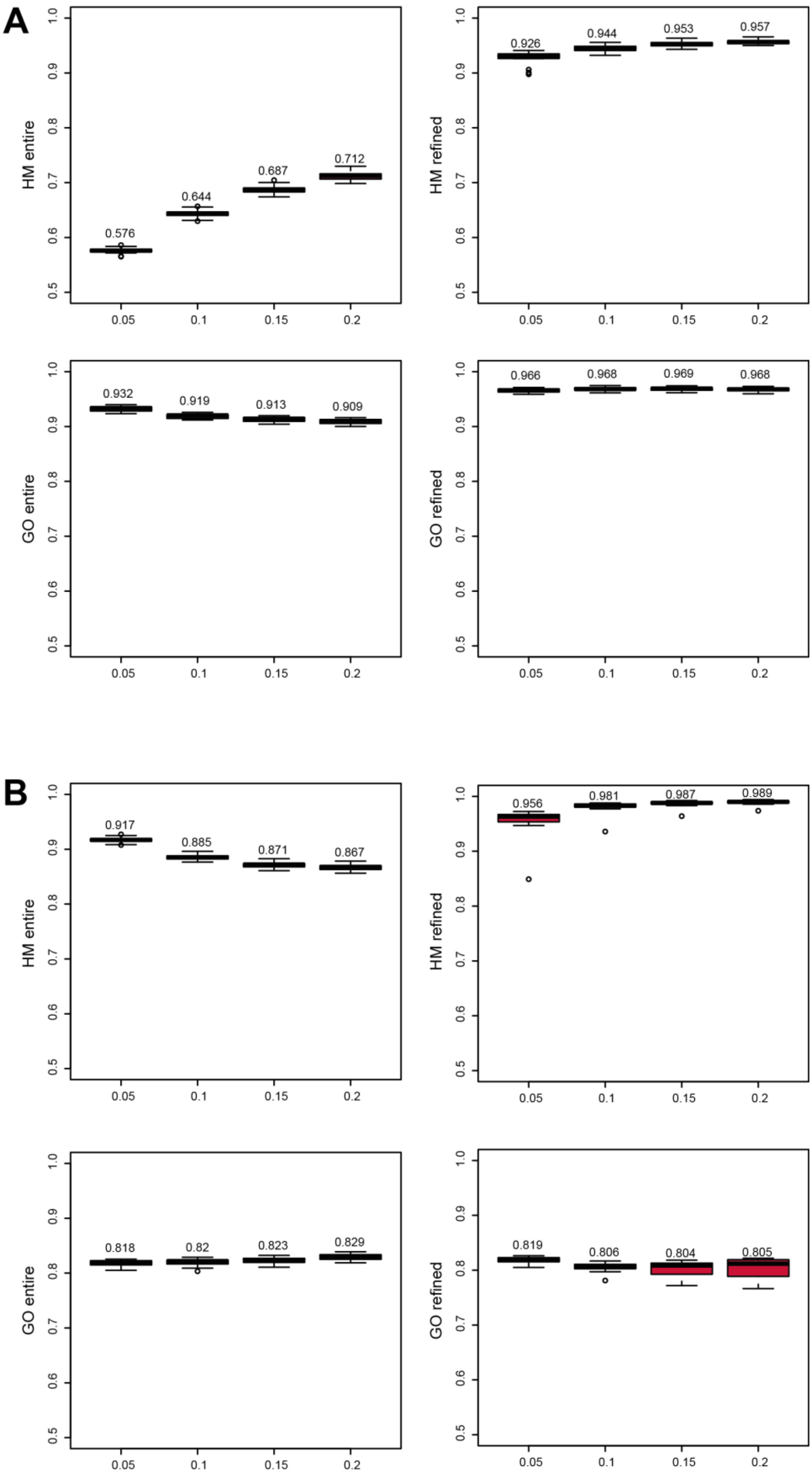


**Supplementary Figure 7** – AUC scores across bootstrap iterations for all A) IFNα response and B) IFNγ response gene sets, using 4 different gene selection thresholds (t). All AUCs were calculated by defining cells post having undergone the induction of interest as the positive control, and the two other populations as the negative control.

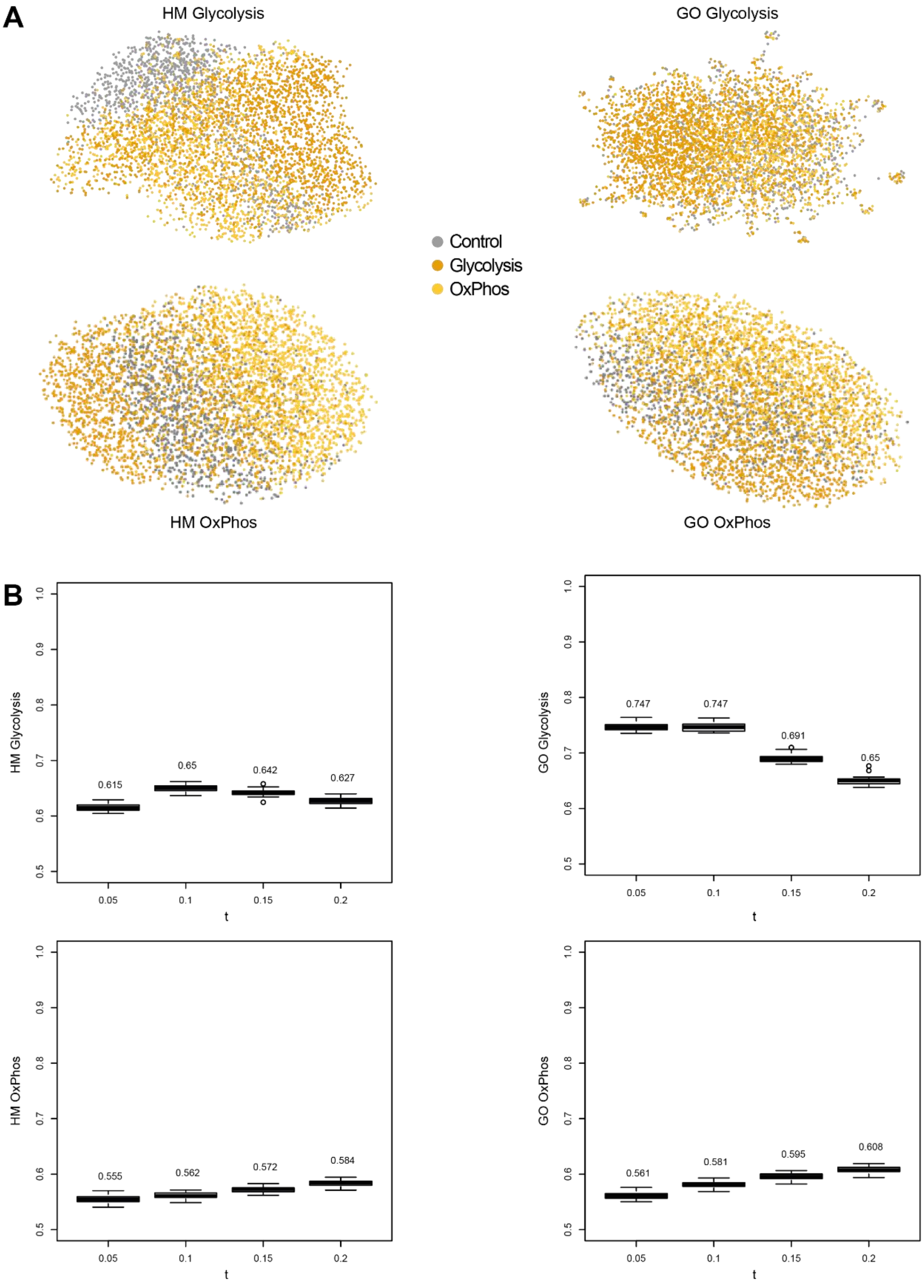


**Supplementary Figure 8** –Performance using original Glycolysis and OxPhos gene sets. A) UMAP population separation for all 3 datasets pooled, using the HM (left) and GO (right) gene sets for Glycolysis (top) and OxPhos (bottom). B) AUC scores across bootstrap iterations for the HM (left) and GO (right) gene sets for Glycolysis (top) and OxPhos (bottom), using 4 different gene selection thresholds (t).

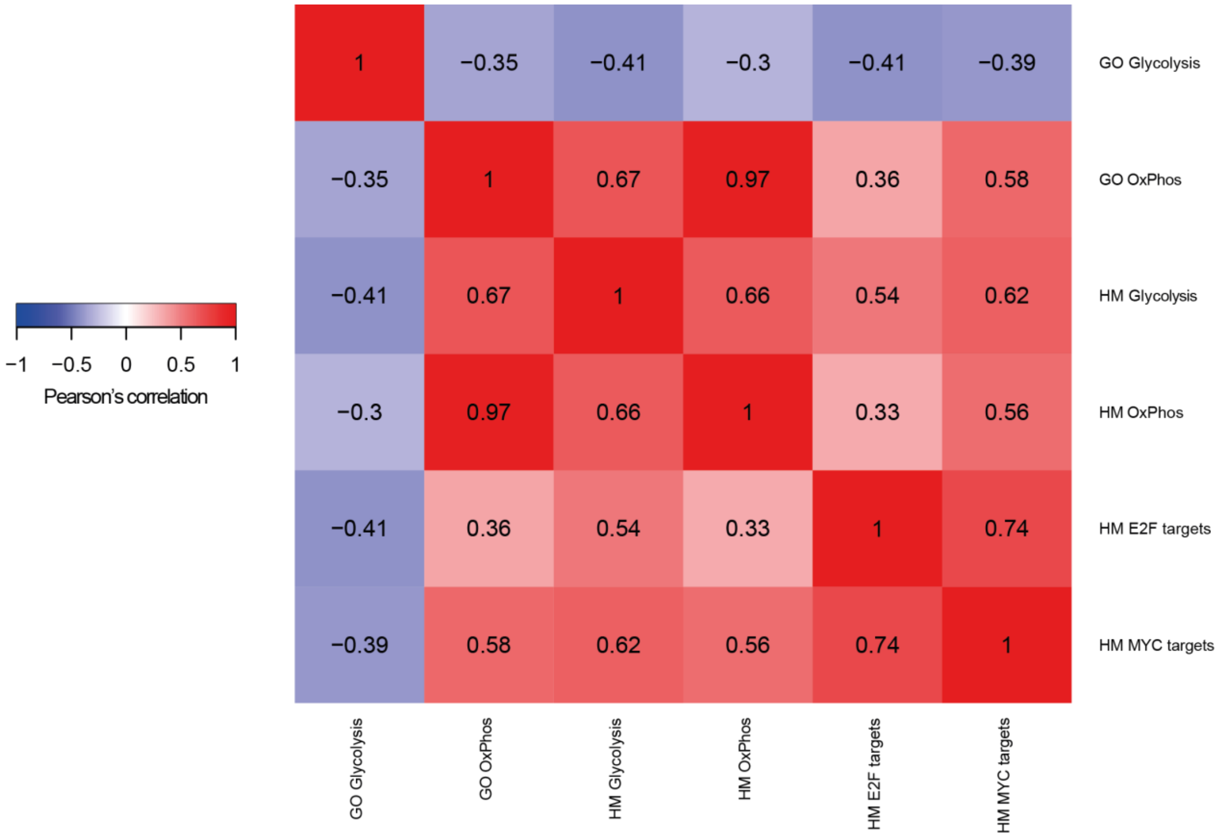


**Supplementary Figure 9** - Correlation heatmap of the glycolysis and OxPhos gene sets from the MsigDB hallmark gene sets (HM) and Gene Ontology (GO), as well as with the proliferation-related HM E2F targets and MYC targets gene sets (0.05 gene selection threshold). Gene set enrichment was performed in the TCGA BRCA dataset. Colors go from blue (perfect anticorrelation) to white (no correlation) to red (perfect correlation).

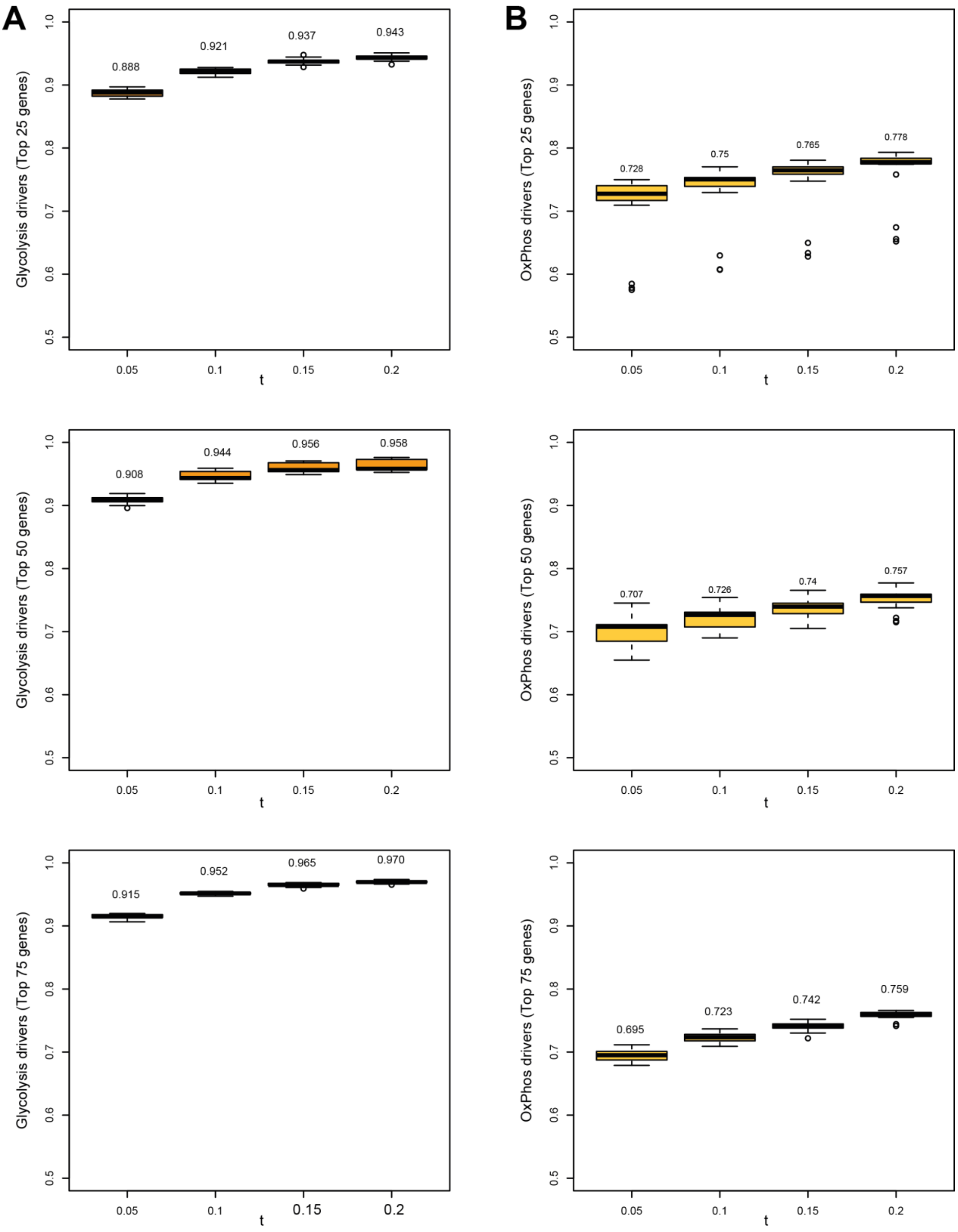


**Supplementary Figure 10** - AUC of the best parameters and gene selection threshold (t) combination for each evaluated glycolysis (A, left) and OxPhos (B, right) gene sets, after validation in external datasets. Combinations of the most significant 25, 50, or 75 genes were separately investigated. All AUCs were calculated by defining cells post having undergone the induction of interest as the positive control, and the two other populations as the negative control.

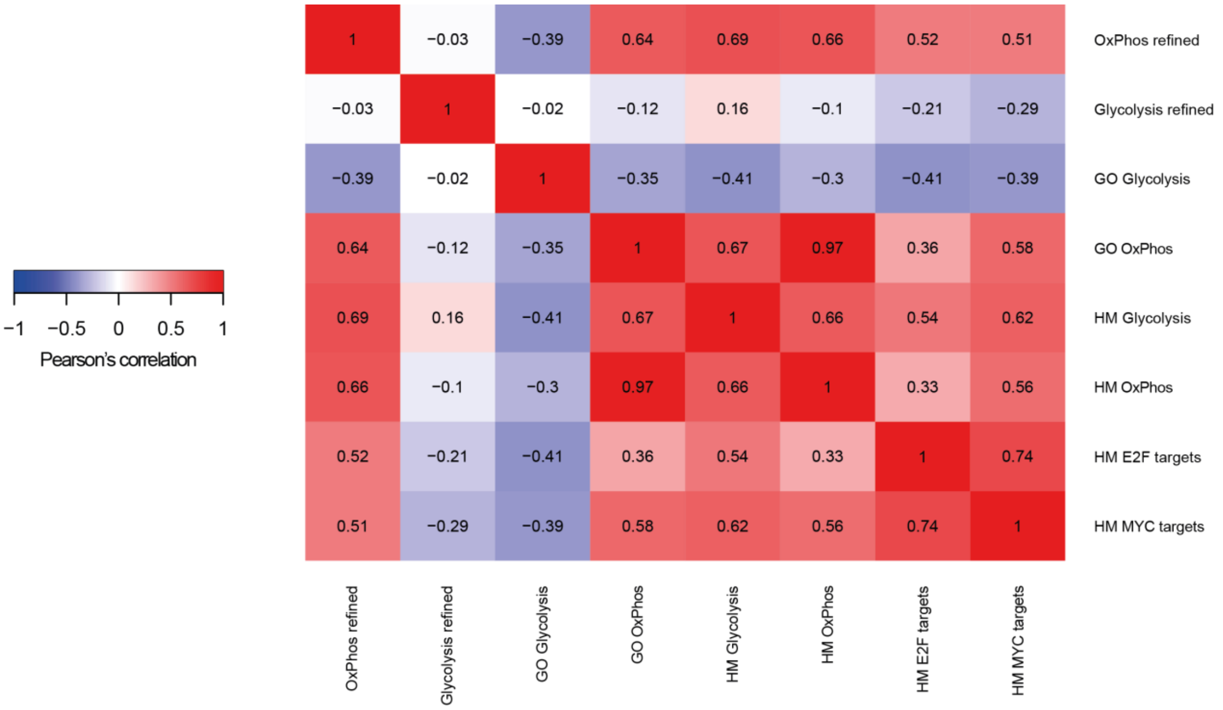


**Supplementary Figure 11** – Correlation heatmap of the refined glycolysis and OxPhos gene sets, the original ones from the MSigDB hallmark gene sets (HM) and Gene Ontology (GO), as well as with the proliferation-related HM E2F targets and MYC targets gene sets (0.05 gene selection threshold). Colors go from blue (perfect anticorrelation) to white (no correlation) to red (perfect correlation).

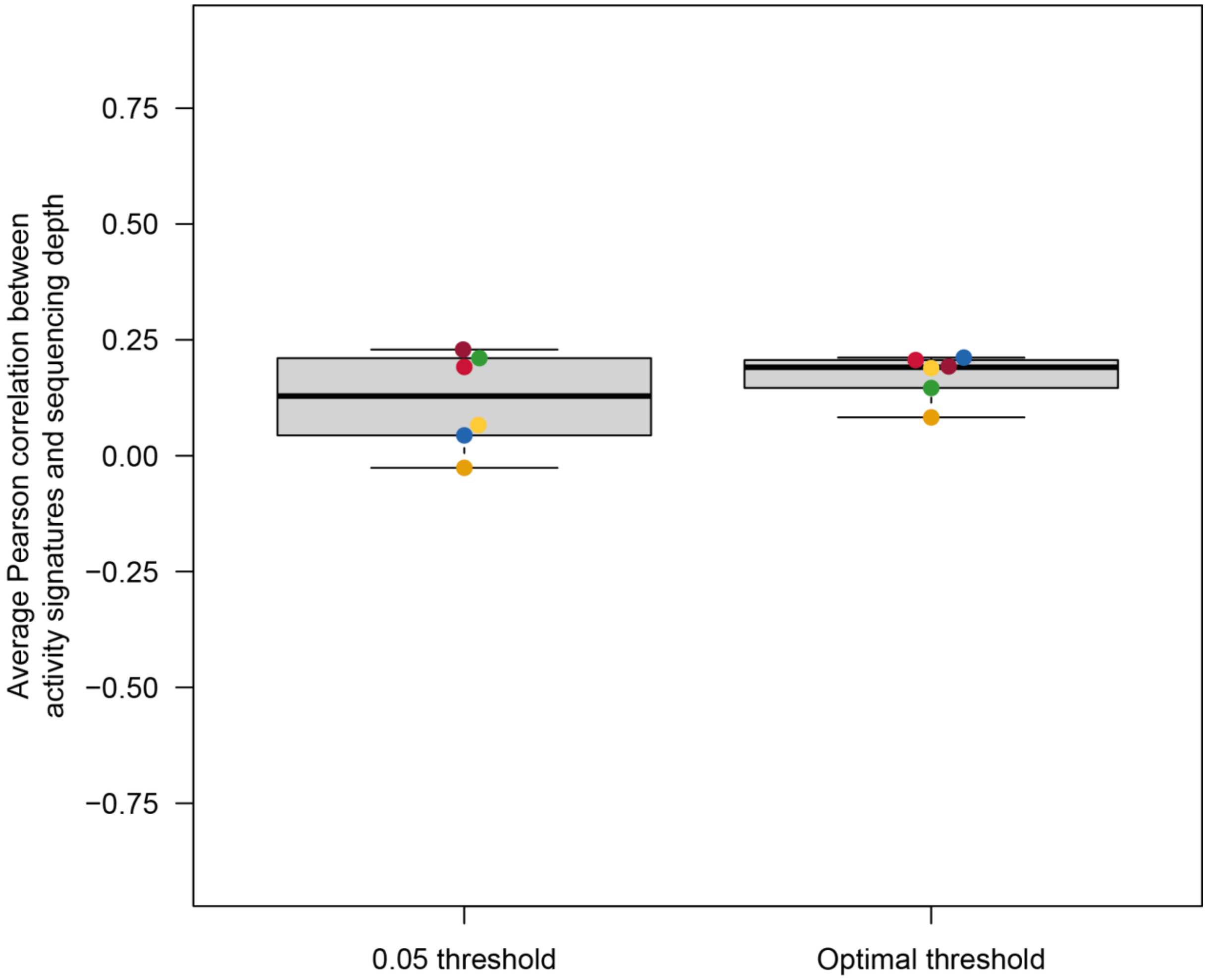


**Supplementary Figure 12** – Correlation between between activity signatures scores and sequencing depth (UMIs per cell). Each dot represents the Pearson correlation coefficient between the two across all cells for a given activity signature: EMT (green), DNA repair (blue), IFNα (dark red) and IFNβ (pink) response, glycolysis (orange) and OxPhos (yellow).

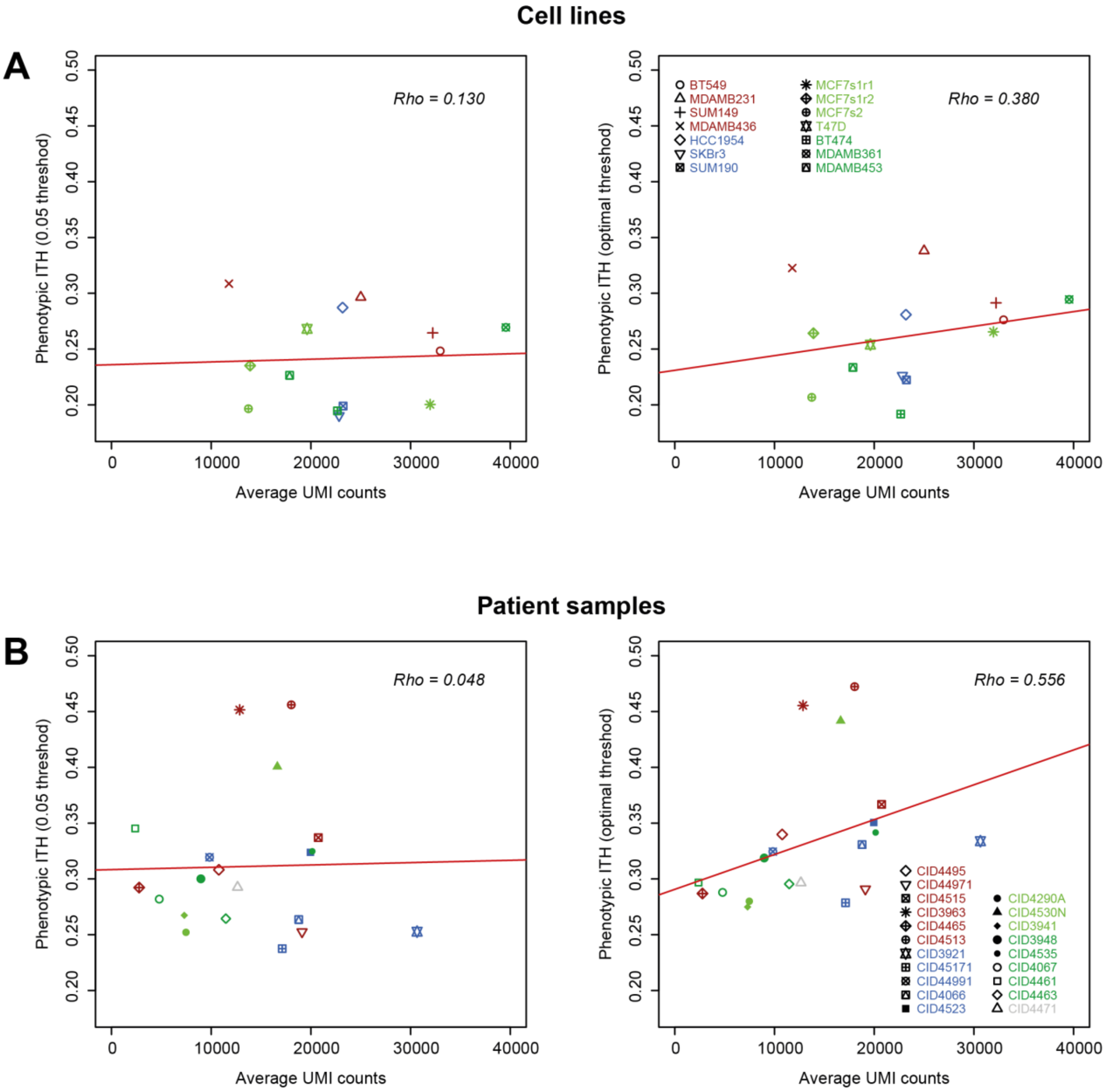


**Supplementary Figure 13** – Correlation between phenotypic ITH and average sequencing depth (UMIs per cell) in A) cell line and B) patient samples, using the recommend 0.05 gene selection threshold (left), or the optimal ones determined using our in vitro induction scRNA-seq data (right). Dots indicate individual samples, color-coded by breast cancer subtype: Basal (red), Her2 (blue), luminal A (light green) and B (dark green), Normal-like (grey). Red lines indicate linear fits.

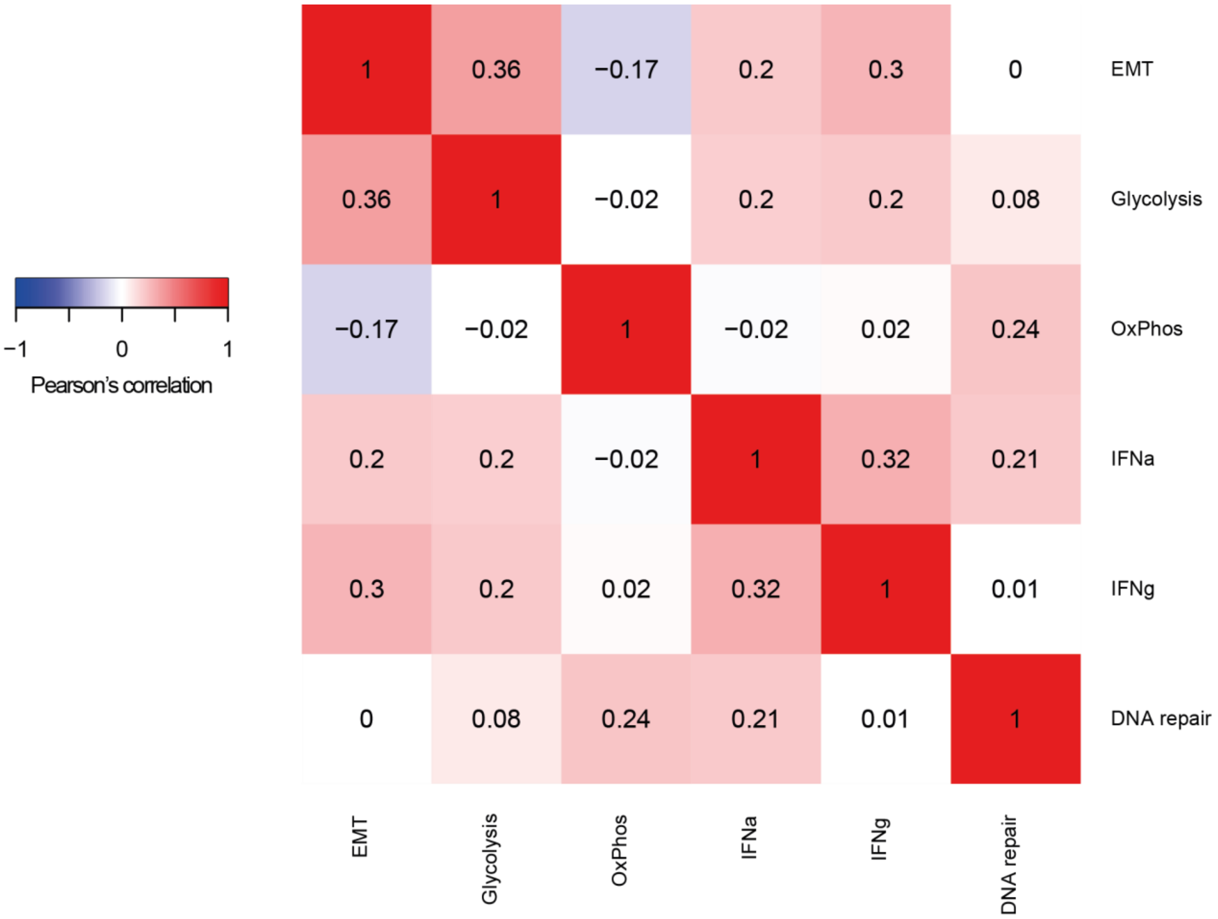


**Supplementary Figure 14** - Correlation heatmap of refined activity signatures using optimal gene selection thresholds. Colors go from blue (perfect anticorrelation) to white (no correlation) to red (perfect correlation).

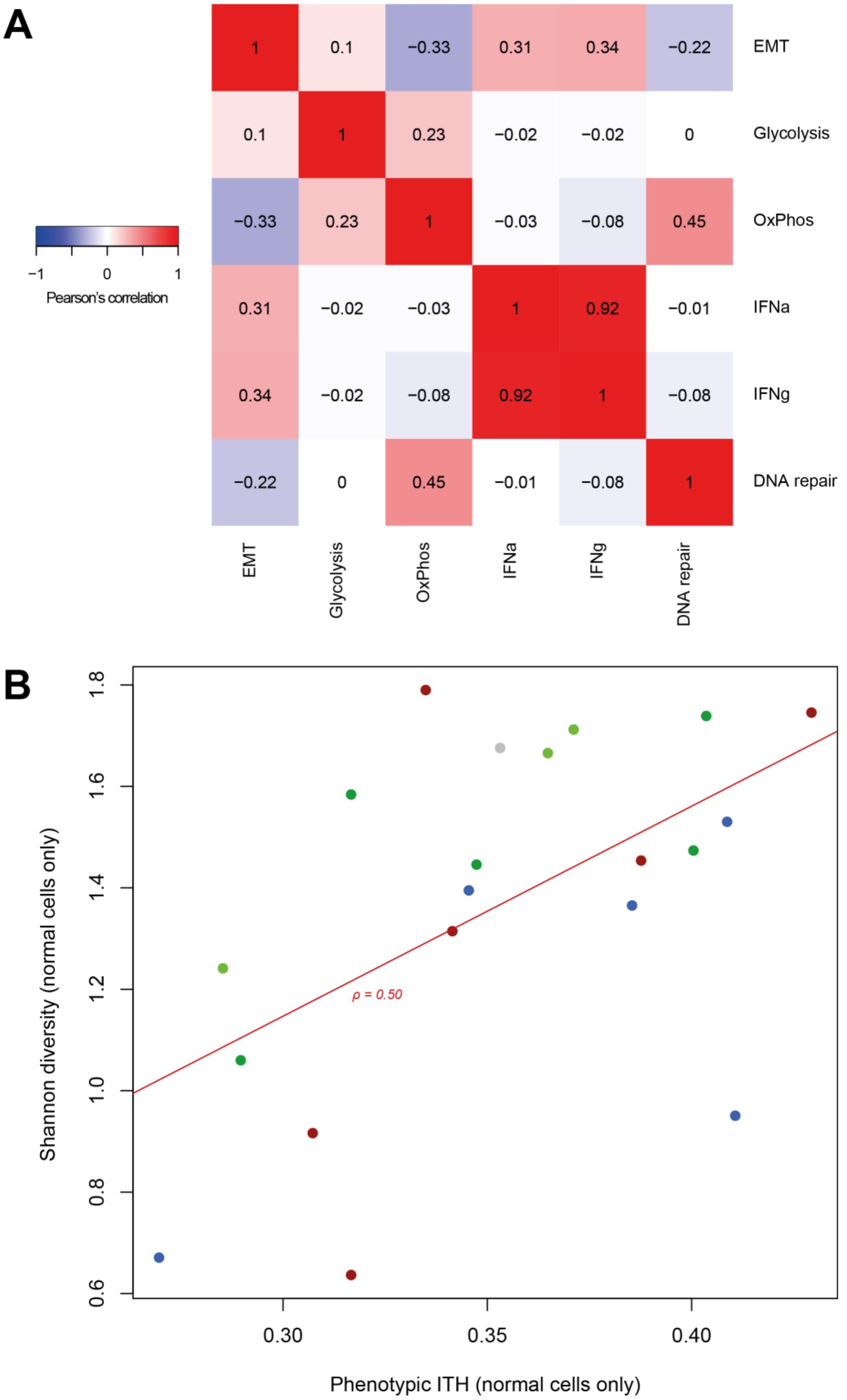


**Supplementary Figure 15** – Performance with entire HM signatures. A) Correlation heatmap of entire HM activity signatures using t=0.05 gene selection threshold. Colors go from blue (perfect anticorrelation) to white (no correlation) to red (perfect correlation). B) Correlation between activity-based ITH, calculated using entire HM signatures and t=0.05, and Shannon diversity index in normal cells from patient samples. Subtypes are color-coded; red line indicates linear regression fit.

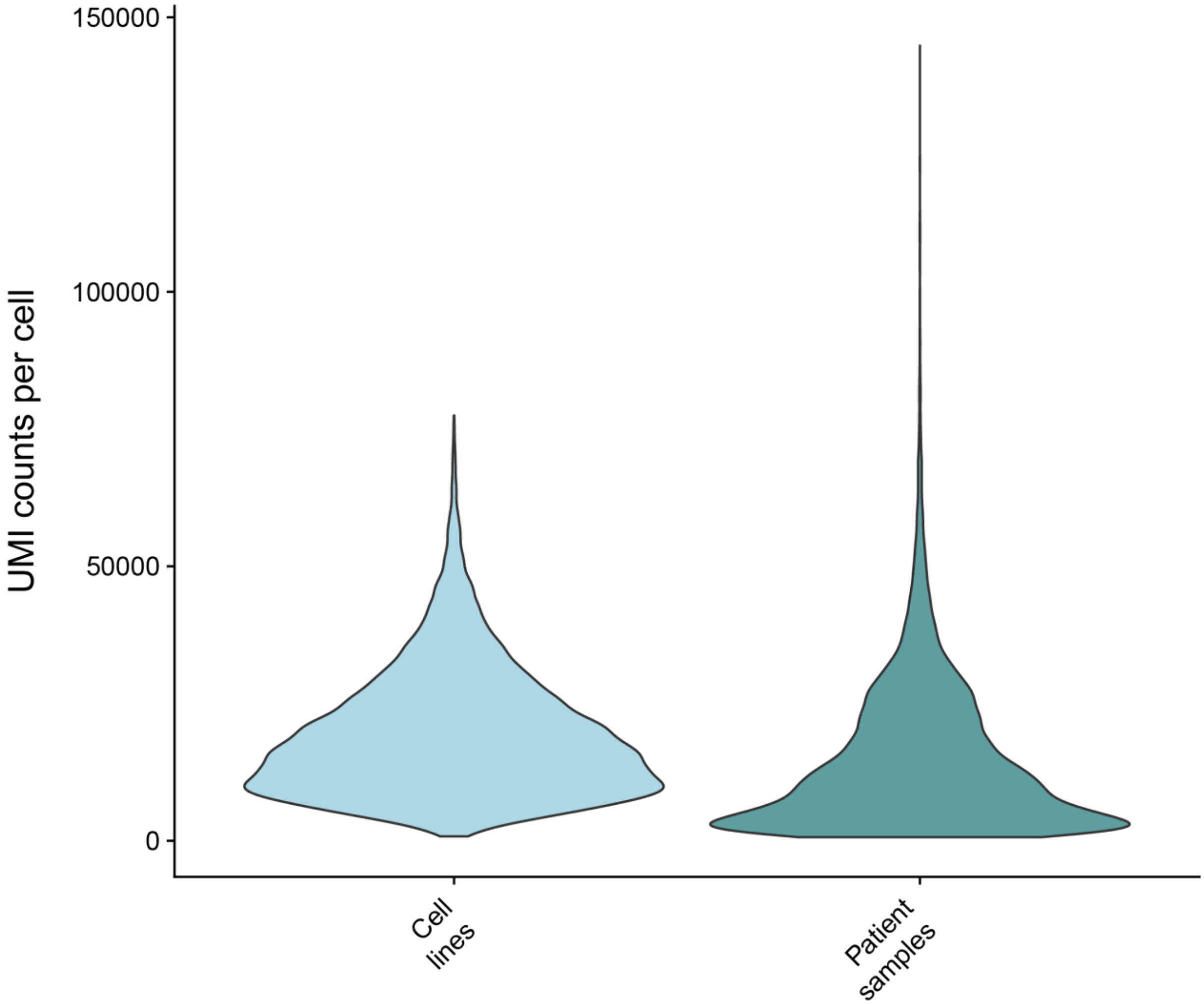


**Supplementary Figure 16** – UMI counts per cell distributions in cell line and patient samples.

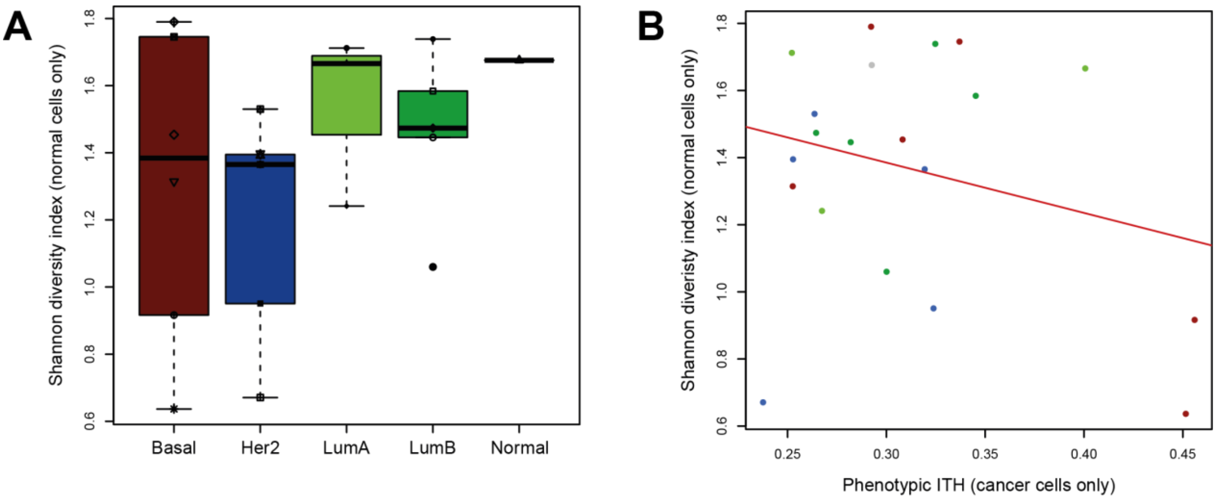


**Supplementary Figure 17** – Classification-based Shannon diversity of normal cells in patient samples. A) Distribution of normal cell Shannon diversity per breast cancer subtype. B) Absence of correlation between a sample's microenvironmental diversity, quantified by Shannon diversity on its normal cells only, and phenotypic ITH, measured only on its cancer cells.

### Supplementary Tables

| | Timepoint | Glucose Concentration (mM) | Protein Concentration (mg/mL) | Normalised glucose consumption |
|---|---|---|---|---|
| Glucose-rich (Glycolysis) | 6h | 6.7 | 3.044 | 0.468 |
| | Overnight | 5.2 | 3.318 | |
| Neutral (Control) | 6h | 6.8 | 2.567 | 0.432 |
| | Overnight | 5.6 | 2.684 | |
| Glucose-deprived (OxPhos) | 6h | 6.9 | 2.284 | 0.404 |
| | Overnight | 5.9 | 2.614 | |

**Supplementary Table 1** - Glucose consumption normalised to population size over time. Protein concentration is used as a surrogate marker of population size.

| **Gene** | **Forward primer** | **Reverse primer** |
|---|---|---|
| AEN | AGGTGGCCCAGAAAGAGATCC | GTGGACATACTTGAGCGCCT |
| PAIMP | CTCACCGTGTGTAGTTGGCA | AGCTGAACACGAACAGTCCT |
| IER5 | AGTTCCCCATTCGTTCTCCG | TGAAGCGGGTCTAGCTTTCC |
| IER3 | GTCTGGTGGTGGGTCGTAAG | CGCCGAAGTCTCACACAGTA |
| XPC | GGAACGAGTTTGGGAATGTG | GTGTAGATTGGGCAGGTTCAG |
| FEN1 | TCTGTGGAGCTGAAGTGGAG | AGTGGATCCCTTGGGTTCTG |
| IRF7 | CCCAGACTGCCTGTGTAGACG | CCAGTCTCCAAACAGCACTCG |
| OAS2 | AGGGAGTGGCCATAGGTGG | AACACCTGGATGGTGAACCC |
| CXCL9 | GAGTGCAAGGAACCCCAGTAGT | TTGTAGGTGGATAGTCCCTTGGTT |
| CXCL10 | TTCAAGGAGTACCTCTCTCTAG | CTGGATTCAGACATCTCTTCTC |
| RPLP0 | GCTGATGGGCAAGAACACCA | CCGGATATGAGGCAGCAGTT |

**Supplementary Table 2** – Primer sequences used for each gene. RPLP0 was used as housekeeping gene for control purposes in all qPCR analyses.

| **Population** | **Cellls before QC** | **Cells after QC** | **N features filter** | **MT genes filter** |
|---|---|---|---|---|
| *EMT* | 1378 | 1198 | >200 | <20% |
| *Ctrl_EMT** | 2379 | 1602 | >200 | <20% |
| *DNA repair* | 875 | 851 | >200 | <20% |
| *Ctrl_DNArep** | 2379 | 1602 | >200 | <20% |
| *IFNα* | 1234 | 1136 | >1000 | <10% |
| *IFNγ* | 1257 | 1151 | >1000 | <10% |
| *Ctrl_IFNs* | 1160 | 1064 | >1000 | <10% |
| *Glycolysis* | 2057 | 1848 | >500 | <5% |
| *OxPhos* | 1649 | 1535 | >500 | <5% |
| *Ctrl_MetabolicPWs* | 1374 | 1226 | >500 | <5% |

**Supplementary Table 3** – QC metrics for in vitro induction scRNA-seq data. *Same control population, used in combination with different post-induction populations.

| Activity | HM gene set | GO gene set |
|---|---|---|
| EMT | EPITHELIAL MESENCHYMAL TRANSITION | EPITHELIAL TO MESENCHYMAL TRANSITION (GO:0001837) |
| DNA repair | DNA REPAIR | DNA REPAIR (GO:0006281) |
| IFNα response | INTERFERON ALPHA RESPONSE | RESPONSE TO TYPE I INTERFERON (GO:0034340) |
| IFNγ response | INTERFERON GAMMA RESPONSE | INTERFERON GAMMA MEDIATED SIGNALING PATHWAY (GO:0060333) |
| Glycolysis | GLYCOLYSIS | POSITIVE REGULATION OF GLUCOSE METABOLIC PROCESS (GO:0010907) |
| OxPhos | OXIDATIVE PHOSPHORYLATION | OXIDATIVE PHOSPHORYLATION (GO:0006119) |

**Supplementary Table 4 –** Gene sets used to detect the presence of each activity, from the MSigDB hallmarks (HM) and Gene Ontology biological process (GO) databases.

| **Activity** | **n_top_genes** | **min_shared_count** | **Seed signature** |
|---|---|---|---|
| EMT | 75 | 25 | HM *EMT* |
| DNA repair | 50 | 50 | HM *DNA Repair* |
| IFNα response | 25 | 50 | GO *IFNα response* |
| IFNγ response | 25 | 50 | HM *IFNγ reponse* |
| Glycolysis | 300 | 75 | GO *Metabolic process* + HM *Glycolysis* + HM *OxPhos* |
| Oxydative Phosphorylation | 300 | 75 | GO *Metabolic process* |

**Supplementary Table 5 –** Optimal parameters for UMAP dimension reduction analyses used to define each activity signature.

| **Breast Cancer Subtype** | **Cell lines** | **Patient samples** |
|---|---|---|
| *Basal* | 4 | 6 |
| *Her2* | 3 | 5 |
| *LumA* | 4 | 3 |
| *LumB* | 3 | 5 |
| *Normal* | 0 | 1 |
| *Total* | n=14 | N=20 |

**Supplementary Table 6 –** Subtype distribution in external breast cancer datasets.

| EMT | DNA repair | IFNa response | IFNg response | Glycolysis | Oxidative Phosphorylation |
|---|---|---|---|---|---|
| LAMC2 | SSRP1 | IFI27 | CXCL11 | ALDH1A3 | GLUL |
| THBS1 | PCNA | ISG15 | CXCL10 | OAT | NAPRT |
| PMEPA1 | TYMS | MX1 | CD74 | RFK | HSD17B12 |
| SERPINE1 | ZWINT | IFIT1 | ICAM1 | PTHLH | CDA |
| TPM4 | XPC | | SOD2 | REXO2 | NIT2 |
| COL4A1 | RRM2B | | TNFAIP3 | MET | ATP5MC3 |
| SLC38A1 | RFC3 | | TNFAIP6 | TGFA | ME1 |
| FN1 | PRIM1 | | | SPTLC2 | GFUS |
| LAMA3 | POLQ | | | TDG | MPC2 |
| ITGA2 | ARL6IP1 | | | GPR87 | FDX1 |
| SFRP1 | DDB2 | | | ALDH18A1 | IRS2 |
| TPM1 | POLR2A | | | AGPS | ADK |
| TAGLN | AK3 | | | NPC2 | ATP5PO |
| FBN2 | POLR1B | | | EPHA2 | DUT |
| DST | SEC61A1 | | | ARPP19 | AIG1 |
| TGM2 | RFC2 | | | PDP1 | VARS1 |
| SERPINE2 | CANT1 | | | PORCN | DBT |
| HTRA1 | POMP | | | SPTLC1 | GOT2 |
| STAG1 | POLR1D | | | PDK3 | ACAT2 |
| TNFAIP3 | POLR3C | | | B3GLCT | CLPX |
| TIMP3 | POLR2H | | | ACOT7 | GLUD1 |
| CALU | ZFP36L2 | | | MTHFD1L | GCLC |
| MCM2 | GTF2H1 | | | CRY1 | NDUFB6 |
| FBLN1 | RAD52 | | | DEGS1 | SCD |
| FGF2 | POLR2E | | | NMNAT2 | ATP5MC1 |
| FLNA | ADA | | | ATP6V1D | ACSS2 |
| VEGFA | TAF13 | | | PRKAA1 | ECHDC3 |
| POSTN | DCTN4 | | | B4GALT4 | NT5E |
| COL12A1 | NCBP2 | | | ICMT | PARK7 |
| GADD45A | AGO4 | | | PDE4D | AIFM2 |
| WNT5A | NELFE | | | PER2 | |
| FERMT2 | POLR2K | | | NBN | |
| DPYSL3 | HPRT1 | | | INPP4B | |
| IL6 | TAF12 | | | EXTL2 | |
| ADAM12 | AK1 | | | PDE8A | |
| FBLN5 | SDCBP | | | GDE1 | |
| SERPINB8 | NME4 | | | EXT2 | |
| MYL9 | NT5C | | | CBR3 | |
| LOX | POLR1C | | | INSIG2 | |
| LRP1 | SUPT4H1 | | | ABCC1 | |
| CYR61 | RBX1 | | | UGGT2 | |
| PTHLH | VPS28 | | | PAPSS1 | |
| DKK1 | GPX4 | | | CARD11 | |
| MFAP5 | ERCC1 | | | ENOPH1 | |
| | MPG | | | MAPK1 | |
| | COX17 | | | MGLL | |
| | MPC2 | | | CELSR1 | |
| | DAD1 | | | PTGS2 | |
| | GUK1 | | | DDAH1 | |
| | CDA | | | GLS | |
| | | | | BLMH | |
| | | | | PGAM1 | |
| | | | | GFPT2 | |
| | | | | PLAAT3 | |
| | | | | EDN1 | |
| | | | | BPGM | |
| | | | | RAB23 | |
| | | | | PDXK | |
| | | | | SAMHD1 | |

| |
|---|
| NAGK |
| NTSR1 |
| PDE7B |
| PODXL |
| PNP |
| FGFR1 |
| CHST12 |
| NOX4 |
| PLA2G4A |
| IGFBP3 |
| LIMA1 |
| PLPP1 |
| SNCA |
| TGFBI |
| ACO2 |
| ATP6V1E1 |
| PGM2L1 |
| LPCAT2 |
| ABCA1 |
| FUT8 |
| COL5A1 |
| STC2 |
| NCEH1 |
| NDUFC1 |
| DSEL |
| TGFB1 |
| CD44 |
| SLC4A7 |
| QSOX1 |
| VCAN |

**Supplementary Table 7** – Gene sets for all 6 final activity signatures.

| | Coefficient Estimate | Std. Error | P-value |
|---|---|---|---|
| Intercept | 0.30182 | 0.01787 | < 2e-16 |
| Subtype (Non-basal) | -0.05571 | 0.01774 | 0.0037 |
| Sampletype (Patient) | 0.07038 | 0.01643 | 0.000165 |

**Supplementary Table 8** – Generalized linear model for the influence of tumor subtype and sample type on phenotypic intra-tumor heterogeneity (without interactions).

| | Coefficient Estimate | Std. Error | P-value |
|---|---|---|---|
| Intercept | 0.30709 | 0.02391 | 1.00E-13 |
| Subtype (Non-basal) | -0.06309 | 0.02829 | 0.0334 |
| Sampletype (Patient) | 0.06159 | 0.03087 | 0.0552 |
| Subtype (Non-basal):Sampletype (Patient) | 0.01241 | 0.03667 | 0.7374 |

**Supplementary Table 9** – generalized linear model for the influence of tumor subtype and sample type on phenotypic intra-tumor heterogeneity (with interactions).
